# The virulence gene *ToxB* is both amplified and disrupted by transposons in the wheat pathogen *Pyrenophora tritici-repentis*

**DOI:** 10.1101/2025.11.13.688278

**Authors:** Ryan Gourlie, Megan C. McDonald, Mohamed Hafez, Reem Aboukhaddour

## Abstract

- Mechanisms that drive virulence gene duplication in plant pathogenic fungi remain poorly understood. In *Pyrenophora tritici-repentis* (*Ptr*), responsible for tan spot of wheat, *ToxB* is a multicopy virulence gene encoding a proteinaceous necrotrophic effector. *ToxB* exhibits a virulence dosage effect, where higher copy numbers are associated with increased disease severity. In this work, we sought to resolve a 25-year old question as to what drove the proliferation of *ToxB* within *Ptr*.
- To investigate this, 23 long-read assemblies were generated and analyzed from a collection of globally distributed isolates with various *ToxB* copy numbers, with a specific focus on regions containing *ToxB*.
- Extensive comparative alignments identified a Helitron-like element, *ToxB-HLE*, that appears to be driving the duplication of *ToxB* in an accessory region of chromosome 4. This region is entirely absent in isolates lacking *ToxB* or its nonfunctional homolog *toxb*. In addition to gene amplification by transposons, multiple independent transposon insertion events were identified in several isolates that disrupted the *ToxB* open reading frame creating inactive *toxb* haplotypes.
- This study provides strong evidence supporting the hypothesis that transposons play dual roles in the rapid evolution of fungal pathogenicity by both amplifying and disrupting a key virulence gene in a globally distributed plant pathogen.

## Introduction

Gene duplications are fundamental drivers of diversification and play a key role in shaping adaptation in eukaryotes by generating genetic redundancy, thereby facilitating the emergence of novel functions (Nei et al., 1969; Holland et al., 1999; Hastings et al., 2009; Ohno et al., 2013; Sacristán et al., 2021). Novel functions emerge when there is relaxed purifying selection on the additional gene copies, while essential functions are maintained by positive selective pressure on the original gene (Kimura, 1983; Long et al., 2003). Gene duplications can occur through many known mechanisms (e.g. retrotransposition, break-fusion-bridge, unequal cross over, transposon capture, etc.), however the impact of these duplications is not always easy to ascertain, particularly for evolutionarily ancient events. The importance of gene duplication in adaptive evolution has been demonstrated in many studies that have shown that ancestral plants, fungi, and vertebrates have undergone multiple rounds of whole genome duplication (Bowers et al. 2003; Kellis et al., 2004; Berthelot et al., 2014). These duplications are often followed by a large reduction in duplicated genes as redundant copies are removed through purifying selection (Kellis et al., 2004; Bloome et al., 2006; Wolf and Koonin, 2013).

Adaptive gene duplications do not need to occur at the whole genome level. In fungi, there are many examples, especially in genes that aid fungi in responding to environmental stress, where particular genes have been duplicated (Wapinski et al., 2007). For example, it has been shown that duplication of biosynthetic gene clusters played an important role in the evolution of trichothecene production in *Fusarium* species (Ward et al., 2002; Brown et al., 2004; Proctor et al., 2018; Rokas et al., 2018). Trichothecenes provide pathogens with a powerful chemical weapon during host infection and confer a strong advantage compared to isolates which lack toxin production. Duplication of single genes rather than whole genomes or gene clusters can also have significant impact. For example, an ancient duplication of the *Sir5* gene, creating *Sir2* in *Zymoseptoria tritici* (a major leaf-spot pathogen), was recently reported to be significantly associated with thermal adaptation (Tralamazza et al., 2024). The authors postulate that the ancient duplication of *Sir2* in a giant transposable element significantly contributed to, and shaped, *Z. tritici*’s ability to survive in highly variable environments (Tralamazza et al., 2024). In *Z. tritici*, the *Sir2* gene remains closely associated with transposable elements that may have driven the duplication event, indicating that these elements can play an important role in initiating gene duplication in fungi.

There are a growing number of examples that demonstrate gene duplications can occur very rapidly under strong selective pressure and remain polymorphic within a fungal population. For example, the duplication of the *CYP51* gene (syn. *ERG11*) decreases sensitivity to azole fungicides (Liu et al., 2011; Stalder et al., 2022; Fan et al., 2023; Arnold et al., 2024). Purifying selection of resistance gene duplications may occur when the high selective pressure of fungicides are removed (Hawkins and Fraaije, 2018). Recently, a virulence gene duplication was explored in the genus *Rhynchosporium* which contains several pathogens of grasses such bromegrass, barley, triticale, and rye. The necrosis inducing effector *NIP1* was found to have substantial copy-number variation in *R. commune*, with higher copy-numbers linked to increased virulence (Mohd-Assaad et al., 2019). In these examples, long-read genome assemblies or large-scale population genomics data were key to uncovering the gene duplication events, because the gene duplications were still highly identical at the nucleotide level. These high-identity copies collapse in short-read assemblies into a single sequence and can be difficult to detect (Mohd-Assaad et al., 2019). This makes determining the exact sequence of events that led to duplications difficult to describe in detail, as these regions often fall into unassembled regions of the genome.

In plant pathogens, transposable elements are drivers of adaptation in regions encoding effectors, small secreted proteins or metabolites that interact and modulate host defense systems (Raffaele et al., 2010; Dong et al., 2015; Torres et al., 2020). In addition, transposons can also alter expression profiles of nearby genes and facilitate genome instability by increasing the likelihood of deleterious rearrangements (Mita and Boeke, 2016; Bhat et al., 2022). Active transposons remain polymorphic in large fungal populations and can thereby drive population specific differences via virulence gene presence/absences and allelic diversity (Fouché et al., 2018; Singh et al., 2021). This variation has a direct impact on the virulence profile of a particular fungal individual in the context of different host genotypes. For example, in the tomato pathogen *Fusarium oxysporum* f.sp *lycopersici* the insertion of a hAT transposon within the effector gene *AVR1* resulted in a loss of function mutation (Inami et al., 2012). While seemingly a negative event, due to the loss of a functional virulence gene, this transposon insertion, under the right host environment, leads to a gain of virulence on host genotypes carrying the *I* resistance gene (Inami et al., 2012). The tension or conflict between transposon activity disrupting important genes versus the adaptive potential of random mutations in fungal pathogen genomes has recently been termed “the devil’s bargain” (Fouché et al., 2022). However, there remains a limited number of examples where transposons have been shown to behave in antagonistic ways on the same fungal gene with a characterised virulence function.

In *P. tritici-repentis* (*Ptr*), causing tan spot of wheat, the effector *ToxB* is one rare example where a known multi-copy virulence gene, that is critically important for disease development, has also been impacted by transposon insertions to create inactive copies (Strelkov et al., 1999; Hafez et al., 2024). ToxB is the second proteinaceous effector identified in the tan spot pathosystem, after the necrosis-inducing effector, ToxA (Orolaza et al., 1995; Lamari et al., 1995; Strelkov et al., 1999). Both effectors function in an inverse gene-for-gene manner, whereby necrosis/chlorosis is only observed in wheat genotypes carrying the corresponding susceptibility genes (*Tsc2* in the case of ToxB) (Lamari and Bernier, 1989; Lamari et al., 1995; Lamari et al., 2003; Faris et al., 2013). *ToxB* was found to be present in field isolates in as many as 10 copies through both PCR techniques and southern hybridization experiments (Martinez et al, 2001; 2004; Strelkov et al., 2005). Through a series of cloning experiments, these *ToxB* duplications were determined to be identical to each other, but the surrounding sequences were different (Martinez et al., 2004). This left a question as to how *ToxB* was duplicated in these genomes.

Importantly, the number of *ToxB* copies has been shown to contribute additively to virulence, leading to quantitative (dosage) effects (Amaike et al. 2008; Aboukhaddour et al., 2012). While these data suggest that the evolution of virulence in this pathogen is driven by duplication of the *ToxB* effector, there also exists a non-functional homolog, designated *toxb,* that is unable to induce chlorosis found in some *Ptr* isolates and other Ascomycetes (Hafez et al., 2024). Recent work found *ToxB* present in 21% of a worldwide *Ptr* population (422 isolates; *ToxB* presence is highly associated with geography) while *toxb* was identified in 6%, indicating that both active and inactive forms persist in the pathogen population, in addition to *toxb*’s presence in other species in the Dothidomycetes (Hafez et al., 2024). The majority of chlorosis-inducing *Ptr* isolates carry the same *ToxB* haplotype (allele), *ToxB1*, though at least three functional *ToxB* (*ToxB1*, *B4*, and *B5*) and five non-functional *toxb* haplotypes (*toxb2*, *b3*, *b12*, *b13*, *b14*, and *b15*) have been identified in *Ptr* (reviewed in Hafez et al., 2024).

Understanding how *ToxB* was duplicated in the tan spot pathogen has been complicated by its multi-copy nature combined with the limited access to *ToxB* coding isolates (Aboukhaddour et al., 2023; Hafez et al., 2024). *ToxB* is mainly found in isolates from the Fertile Crescent, absent from Australian isolates and very rare in isolate collections from North America. In this study, a novel collection of *ToxB* containing isolates obtained from the Middle East offered a unique opportunity to further explore the evolution of this important effector. Our previous work using long read assemblies of isolates I-73-1 and D308 showed that *ToxB* (and its homolog *toxb*) were carried on chromosome 4 (Chr04) within what appeared to be an accessory region (Gourlie et al., 2022). This region shared several common features with a *Starship* transposon, but was missing the tyrosine recombinase captain, and hence it was thought it may be an ancient/derelict *Starship* transposon (Gourlie et al., 2022). Earlier research using southern hybridisation approaches indicated that *ToxB* can be carried on two different chromosomes in the same isolate (Alg4-X1) (Aboukhaddour et al., 2009). Recently, a Copia-like retrotransposon (*Copia-1_Ptr*) was identified in some isolates of *Ptr* which has disrupted the open-reading frame of *ToxB*, resulting in an inactive form of *toxb* (Hafez et al., 2024). Together, these studies suggest that transposon activity may be a key driver in both the expansion and inactivation of the *ToxB* gene in different populations. To address these questions, 23 long-read assemblies of *Ptr* from seven countries were generated, including eight isolates with *ToxB*, nine with the inactive *toxb* allele, and six isolates completely lacking either *ToxB* or *toxb* based on previous PCR and southern blot hybridization analyses (Aboukhaddour et al., 2009; 2013; Kamel et al., 2019; Wei et al., 2021; Hafez et al., 2024). This comprehensive collection of diverse isolates provides a framework to dissect the genomic mechanisms that led to *ToxB* duplication and pseudogenization.

## Methods

### Isolates, DNA extraction, and sequencing

In total, 24 isolates were included in this study, details presented in Table 1. The isolates span 65 years from North America, Europe, Japan, North Africa, and the Fertile Crescent. These isolates have been studied by number of labs over the past 30 years, have advanced research on fungal virulence evolution and plant-pathogen interaction. Many of these isolates have been included in previous works and are well studied. DNA was extracted as described in Gourlie et al., (2022). Briefly, isolates were derived from a single spore culture as described in Aboukhaddour et al., (2013) and grown in 100 mL PDB flasks @ 25°C in the dark. Mycelia mats were harvested after ∼2 weeks then washed several times with milli-Q water and freeze-dried as described in Gourlie et al., (2022). Genomic DNA extracted using Genomic Tip 100/G kits from Qiagen. A subset of isolates were sequenced in 2022 by Genome Quebec using PacBio Sequel, and the remaining isolates were sequenced at Mayo Clinic in 2023 using PacBio RS II (sequencing details provided in Methods S1).

**Table 1.**
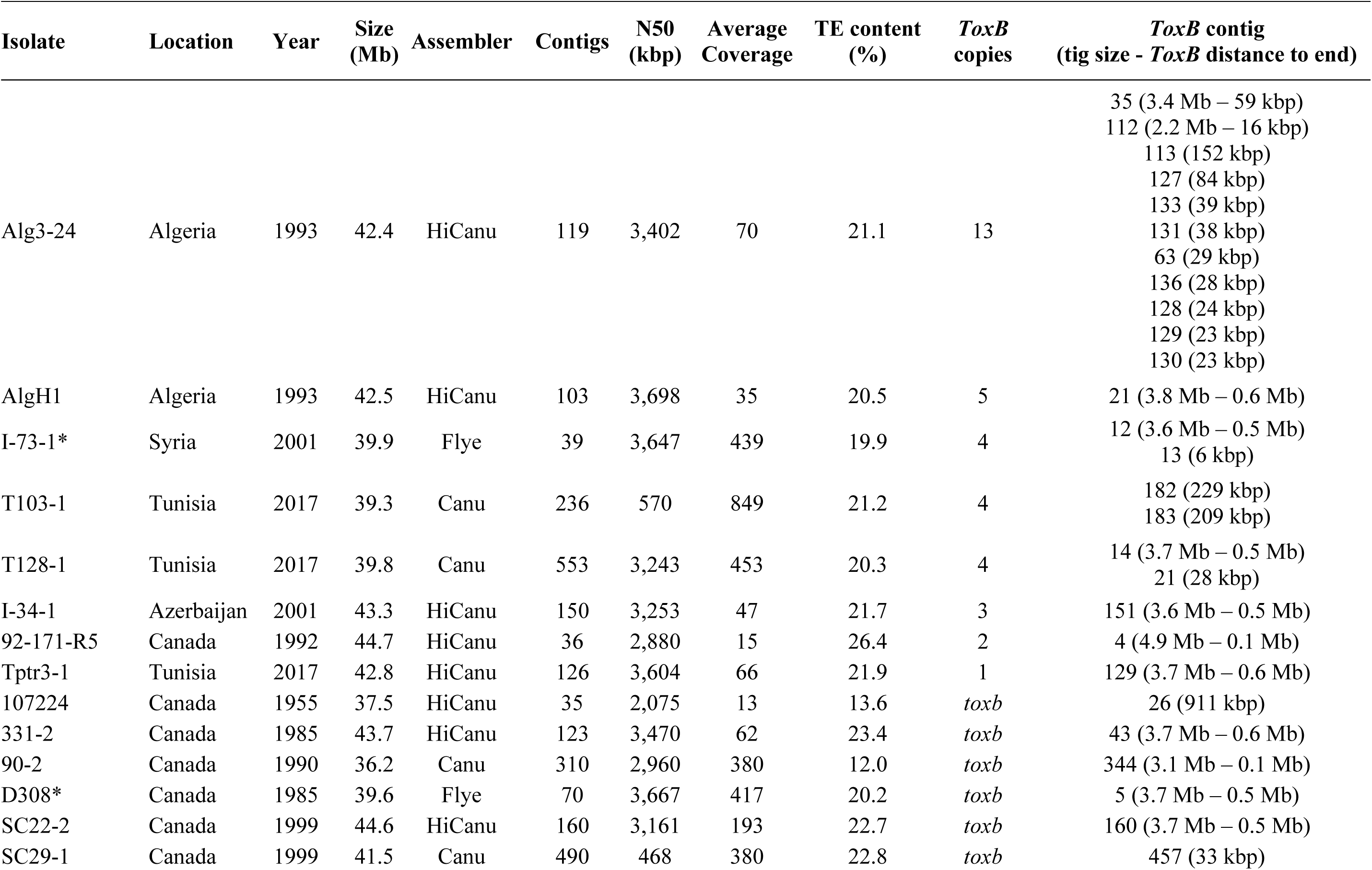
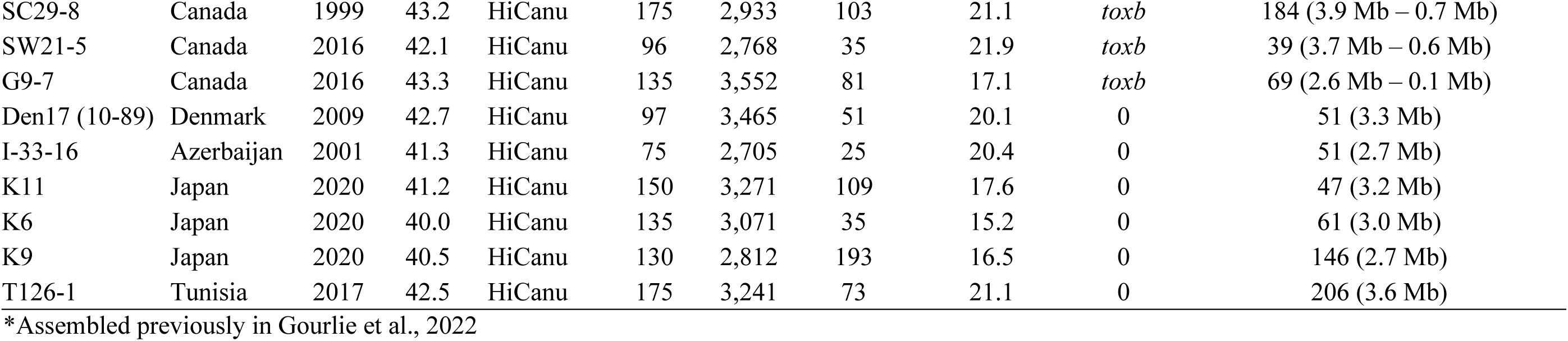
Assembly statistics and *ToxB* information of *Pyrenophora tritici-repentis* genomes used in this study. Table is ordered by *ToxB* copy number, followed by isolates with *toxb*, then isolates without *ToxB* or *toxb* (alphabetical by isolate name within groupings). Genomic data available in BioProject PRJNA803191.

### Genome assembly

The subset of isolates sequenced at Genome Quebec were assembled using CANU v2.2 after converting subread.bam files to fastq using SAMtools v1.9 (Li et al., 2009; Koren et al., 2017). The set sequenced at Mayo Clinic were assembled using HiCanu v2.2 (Nurk et al., 2020). For both CANU and HiCanu the default settings were used. QUAST (Gurevich et al., 2013) and BUSCO scores were used to assess assembly quality using the Pleosporales db10 dataset (Simão et al., 2015). Completeness and *ToxB* regional confidence was also assessed by aligning raw reads to the assemblies using minimap2 v2.18-r1015 and SAMtools v1.9 (Li et al., 2009, Li, 2018). Total transposon content of each isolate was assessed with EDTA (v2.2) and/or EarlGrey v4.2.4 (Ou et al., 2019; Baril et al., 2024). A basic linear regression model was fit to the assembly size by percentage of base pairs annotated in EDTA to asses if there was a relationship.

### Alignments

Alignment and sorting of reads to assemblies was done with Bowtie2 (Langmead and Salzberg, 2012) and SAMtools v1.9 (Li et al., 2009) then visualized with either IGV (Robinson et al., 2011) or dotplotly (Porten, 2018). Whole genome alignments were done with minimap2 (v2.18-r1015) (Li, 2018) and visualized with dotplotly (Porten, 2018). Genome alignment also done with progressiveMauve (Darling et al., 2010). Regional/local alignments were done within Geneious built-in dotplotter (Biomatters Ltd.), MAFFT v1.5.0 (Katoh and Standley, 2013), LASTZ v7.0.3 (Haris, 2007). Chromosome scale alignments were visualised with the gggenomes R package (Hackl et al., 2024). Circular alignments were done with Sibelia v3.0.7 (Minkin et al., 2013) and visualized with Circos (v0.69-8) (Krzywinski et al., 2009). Reference genomes Pt-1C-BFP (GCF_000149985.1) and M4_2.2 (GCA_003171515.3) were retrieved from NCBI for alignments (Manning et al., 2013; Moolhuijzen et al., 2018).

### Identification of the ToxB Helitron-like-element (ToxB-HLE) and other transposons

Each copy of *ToxB* from each genome including various amounts of upstream and/or downstream sequence were extracted. The up/down stream sequences ranged between 50 bp to 15 kbp, stopping at the next *ToxB* copy when in tandem. Multiple iterations of alignments (using tools described previously) were used to identify where sequence similarity ended between copies to establish putative edges/edge sequences. Putative edge sequences were used to extract copies of the repetitive region containing *ToxB*. Open-reading frames were identified within and just beyond the putative edges using Geneious ORF finder with default settings. ORFs were used to BLAST the UniProt database (Consortium, 2015) to provide functional information or identify putative domains. The shortest edge-to-edge sequence was used as a reference to compare copies. Variations were manually catalogued. Hairpin structures were identified using VectorBuilder v2.1.813. When comparing different copies of the *ToxB* region polymorphic insertions >50 bp were further explored using similar methods: identifying edges, determining edge type (i.e. LTR or TIR), search for target site duplications, ORF prediction and annotation, BLASTN of RepBase (vSep2020), and literature comparison. Transposon schematics were created manually. To determine the association of specific DNA sequences with mobile elements, BLASTN hits (90% ID over 90% query) of sequences of interest were intersected with output from EDTA using bedtools. Annotations from EDTA containing the sequences were extracted and aligned using MUSCLE v5.1 (Edgar, 2004). A maximum likelihood phylogeny was constructed from the alignment using the Tamura-Nei model with 50 bootstrap replicates in MEGA-X (Kumar et al., 2018).

## Results

### Long-read assemblies reveal genome expansion is driven by transposons

The isolates selected for sequencing represent diverse geographical regions from North America, Japan, North Africa, and Caucasus region with sampling times that spanned from 1955 to 2020 (Table 1). Isolates were selected based on their estimated *ToxB* copy number, along with a subset known to carry the inactive homolog, *toxb.* Six isolates that completely lack *ToxB* or *toxb* as determined previously were also selected (Kamel et al., 2019; Hafez et al., 2022). All isolates were *de novo* assembled with CANU or Hi-CANU (depending on sequencing technology used) with an expected genome size of ∼40 Mb based on the reference isolate M4_2.2. Assembled genome sizes varied from 34.5 to 44.7 Mb (Table 1). Genomes assembled from HiFi data (circular consensus) contained more contiguous sequences than genomes assembled from older continuous long-read sequencing data (i.e. fewer and larger contigs; Table 1). Genome completeness was assessed using BUSCO and all assemblies had greater than 96% BUSCO scores based on the Pleosporales database (pleo_db10). The transposon annotation pipeline EDTA identified transposon content across isolates ranging from 12 to 26% of the total genome (Table 1). Genome size increased linearly with total predicted transposon content (R^2^ = 0.58; p<0.001) (Fig. 1). The reported R^2^ value suggests transposons are responsible for 58% of the observed genome expansion with some other unknown variable(s) contributing the remaining ∼42%. The main drivers of transposon expansion are LTR retrotransposons (i.e. Copia, Ty3, and unknown elements with LTRs) as well as DNA hAT transposons.

**Fig. 1.**
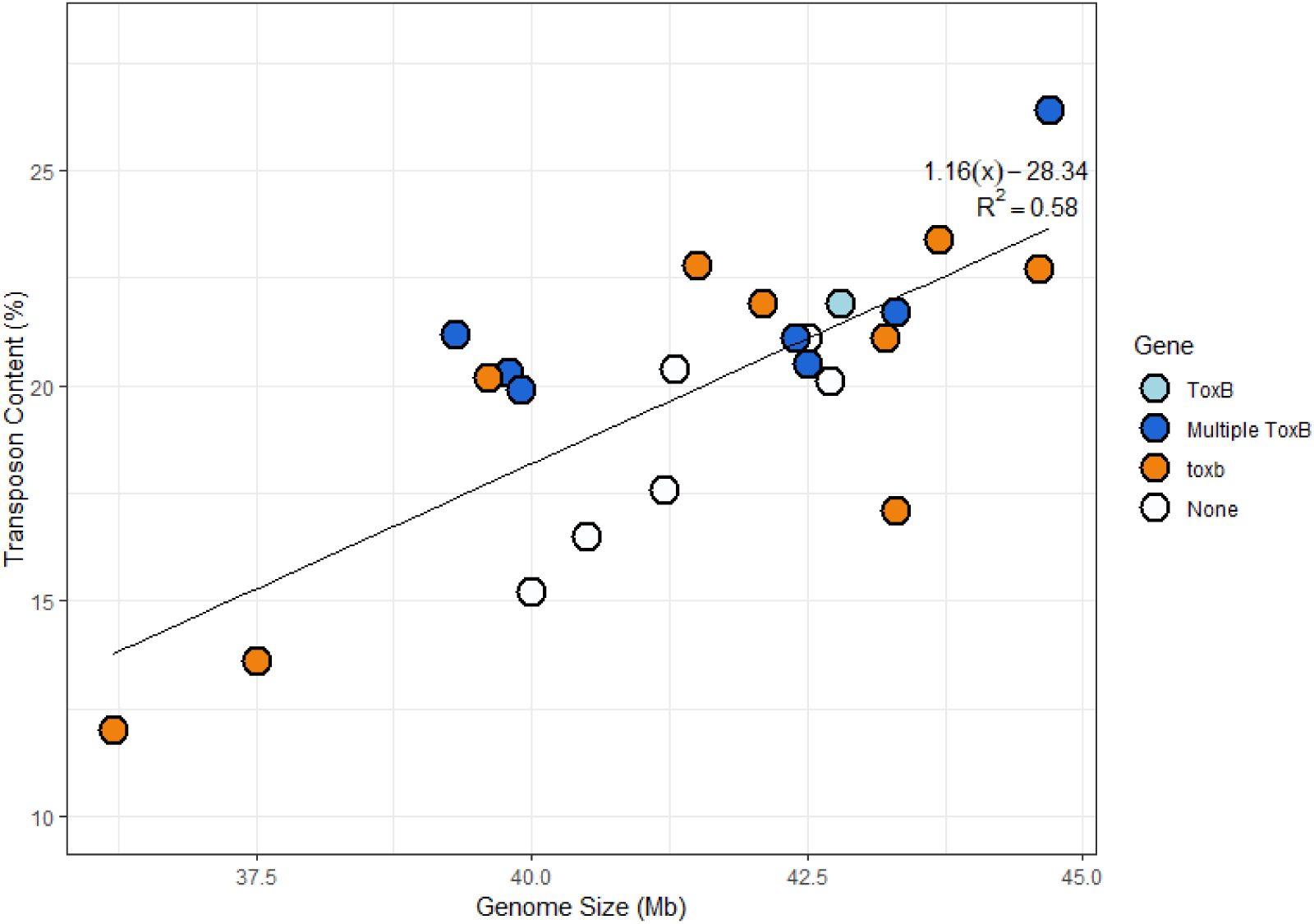
Comparison of genome size versus total transposon content predicted by EDTA. Each dot represents an individual isolate and the dot color indicates the *ToxB/toxb* genotype as indicated by the legend. A linear model fit to the data with an R^2^ value of 0.58 indicates approximately 58% of the size variation could be attributed to transposon content, with the remaining 42% can be attributed to other causes.

### Whole chromosome alignment shows accessory chromosome arm associated with *ToxB* presence/absence

BLASTN of the *ToxB* open reading frame (ORF) revealed *ToxB* copy-numbers in each isolate, their chromosomal location and their relative positions to other copies (Fig. 2). As expected, based on previous PCR genotyping and phenotypic screens six isolates, Den17, T126-1, K11, K9, K6, and I-33-16, did not carry *ToxB/toxb* (Kamel et al., 2019; Hafez et al., 2022; Aboukhaddour unpublished data). The functional *ToxB* was found as multiple copies in seven isolates: Alg3-24, AlgH1, I-34-1, I-73-1, T103-1, 92-171-R5, and T128-1. Isolate Tptr3-1 contained only a single functional copy of *ToxB*. Nine isolates, 90-2, 331-2, D308, G9-7, SC22-2, SC29-1, SC29-8, SW21-5, and 107224, carried a single non-functional *toxb*. No isolates contained multiple *toxb* copies, and no isolate contained a combination of *ToxB* and *toxb* together. Of the 17 *ToxB/toxb* containing isolates, 13 assemblies harbour the gene on chromosome sized contigs (defined here as ≥1 Mb). The remaining four assemblies contained *ToxB/toxb* on contigs ranging from 300 kbp down to only 27 kbp (Table 1). Based on alignments with the reference isolates Pt-1C-BFP (non-carrier; USA) and M4_2.2 (non-carrier; Australia) *ToxB/toxb* is located on Chr04 in 12 isolates. Three isolates, Alg3-24, 92-171-R5, and G97, had *ToxB* copies assembled into different chromosomes. In the Algerian isolate, Alg3-24, one *ToxB* copy was found on Chr05 and an additional two copies were found on Chr10 (Fig. 2). In addition, Alg3-24 had 10 *ToxB* copies on contigs ≤ 150 kbp. It is not clear if these smaller contigs represent additional copies in other chromosomes and attempts to manually align these smaller contigs yielded inconclusive results. As such they were omitted from further analysis. Isolates 92-171-R5 and G9-7 are phylogenetically distant from the other *Ptr* isolates (Gourlie et al., 2022) and *ToxB/toxb* was found on Chr03 and Chr07 respectively, rather than chromosome 4 as with most isolates. Finally for two isolates, SC29-1 and T103-1, *ToxB/toxb* assembled into smaller contigs (∼30 kbp and ∼200 kbp) where we could not determine its precise location in the genome as they only aligned to an accessory region (described in more detail below). Chromosome alignment of Chr04 *ToxB/toxb* isolates with isolates that do not carry *ToxB/toxb* revealed a large accessory region (Fig. 3). When aligned to the triple copy *ToxB* isolate I-73-1, the gap size in non-*ToxB* isolates ranged between ∼50 kbp to ∼375 kbp (Fig. S1). In comparison, this region was largely syntenic in alignments between *ToxB* and *toxb* isolates, indicating a large insertion in isolates carrying *ToxB/toxb* (Fig. 3).

**Fig. 2.**
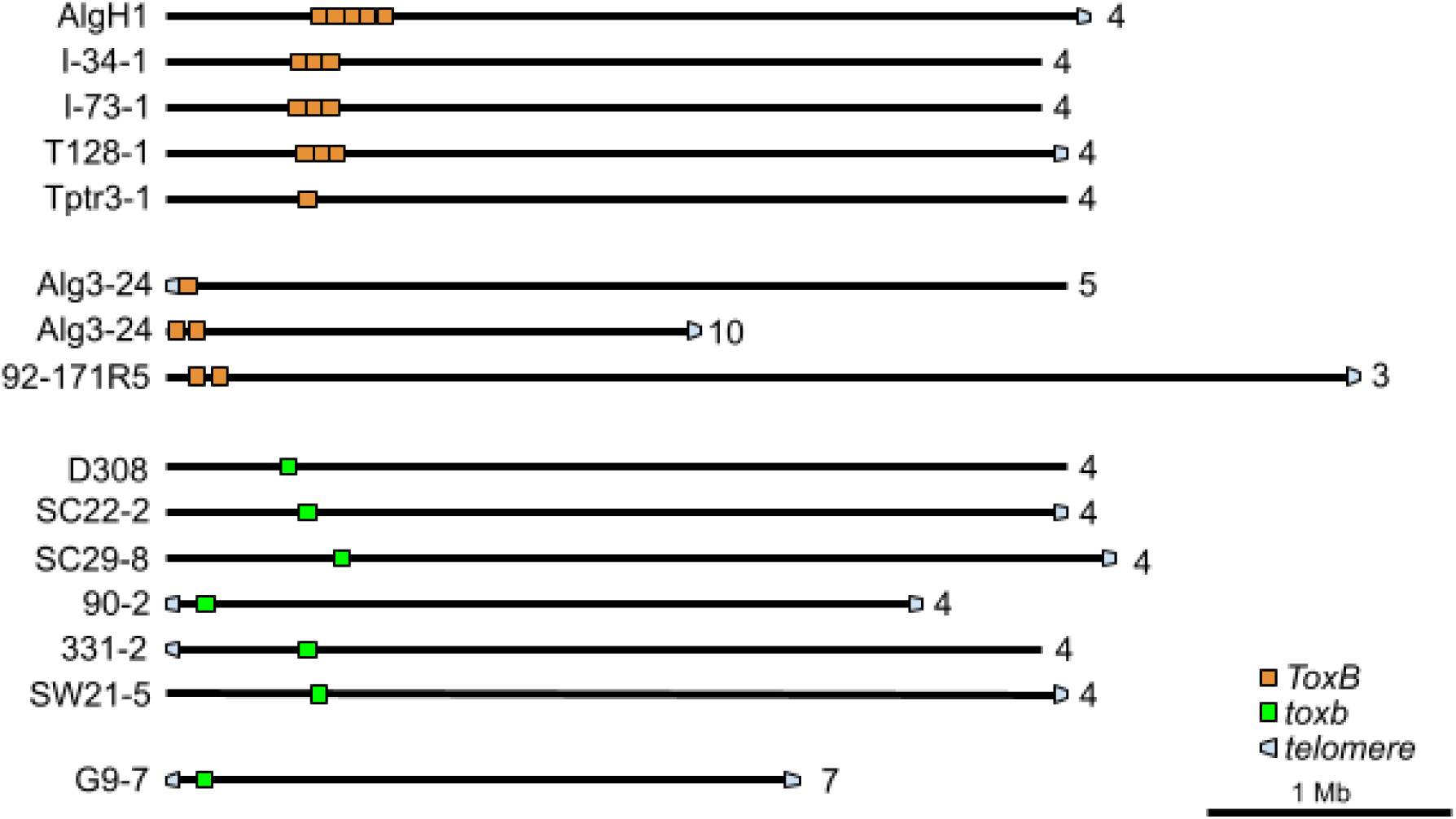
Relative positions of *ToxB/toxb* copies in isolates with chromosome sized contigs (>1 Mb). Orange boxes indicate active *ToxB* isoforms and green boxes indicate inactive *toxb* haplotypes. Grey caps indicate presence of telomeric repeats (TTAGGG/CCCTAA). The number on the far right represents the corresponding chromosome number when aligned with M4_2.2.

**Fig. 3.**
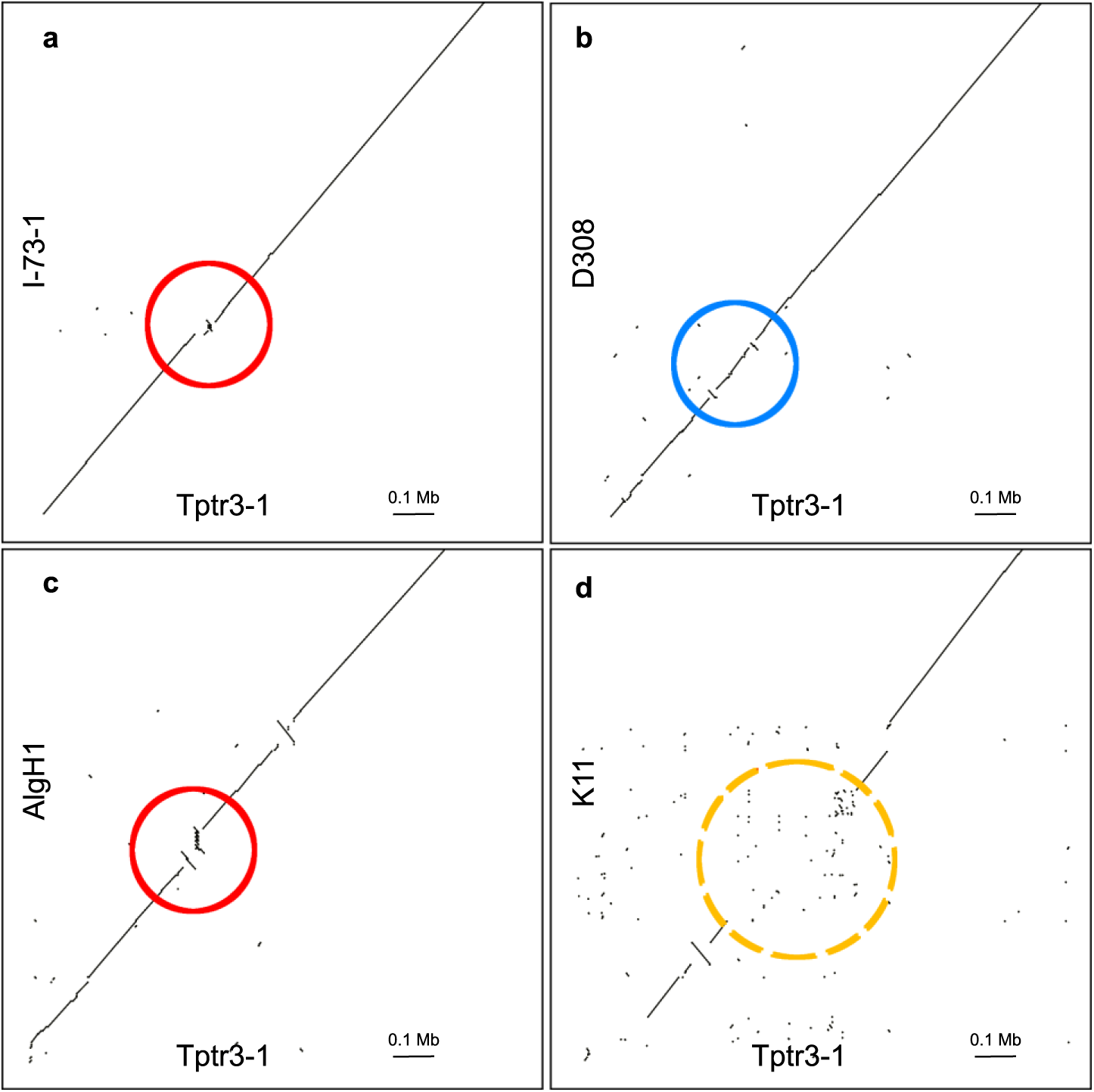
Representative dotpot alignments of *ToxB/toxb* containing and non-containing contigs. The x-axis in all alignments is a section of Chr04 from isolate Tptr3-1 which contains a single copy of *ToxB.* The y-axis in plots A-D are the same section of Chr04 from different isolates: **a** alignment with I-73-1 which contains three *ToxB* copies. Area containing *ToxB* is highlighted in the red circle; **b** alignment with *toxb* isolate D308, region containing *toxB* is highlighted in a blue circle; **c** alignment with isolate AlgH1, which contains five copies of *ToxB* marked by a red circle; and **d** alignment of K11 that does not contain *ToxB* or *toxb*. The accessory region is marked in an orange dashed circle.

### *ToxB* copies are present as tandem duplications

The BLASTN results reported above identified five isolates where multiple copies of *ToxB* were localized to the accessory region of Chr04 in close proximity to each other. Closer inspection of this region showed that *ToxB* and some flanking DNA were arranged in tandem copies (or direct repeats). In AlgH1, we identified five tandem copies, while in I-34-1, I-73-1, T103-1, and T128-1 we identified three tandem copies (Fig. 2 and 4). To assess whether this tandem array of *ToxB* was not due to a mis-assembly of this region, the raw PacBio reads were aligned against the *de novo* assembly for each isolate. Isolates with three tandem *ToxB* copies (I-34-1, I-73-1, T103-1, and T128-1) all have single reads which span the entirety of the *ToxB* triplication (Fig. S2), providing high confidence that the tandem array of *ToxB* genes were assembled correctly. For AlgH1, with five tandem copies, we identified five reads with two of the five tandem copies present but no single read that spanned the full 56 kbp region. Coverage of the *ToxB* region is reasonably high (30x). In Alg3-24, we did not observe the same tandem copy structure seen in the previously listed isolates, rather two copies with inverted orientation were found on a 3.4 Mb contig that was homologous with Chr05 and a single copy on a separate 2.2 Mb contig homologous with Chr10 (Fig. 2; Table 1). In this assembly, there were many other *ToxB* copies on small contigs (∼20-80 kb), all with high transposon content (not shown) likely explaining their inability to be assembled contiguously despite the high average genome coverage (∼70x). All *ToxB* containing reads from Alg3-24 contained only single copies of *ToxB*, therefore it is difficult to conclude if the assembly showing two copies on one contig is correct (Fig. S3). Using the same method of aligning raw reads back to the assembly, we also validated the *de novo* assemblies of seven isolates with a single inactive *toxb*, and six assemblies which lack *ToxB* or *toxb* entirely.

**Fig. 4.**
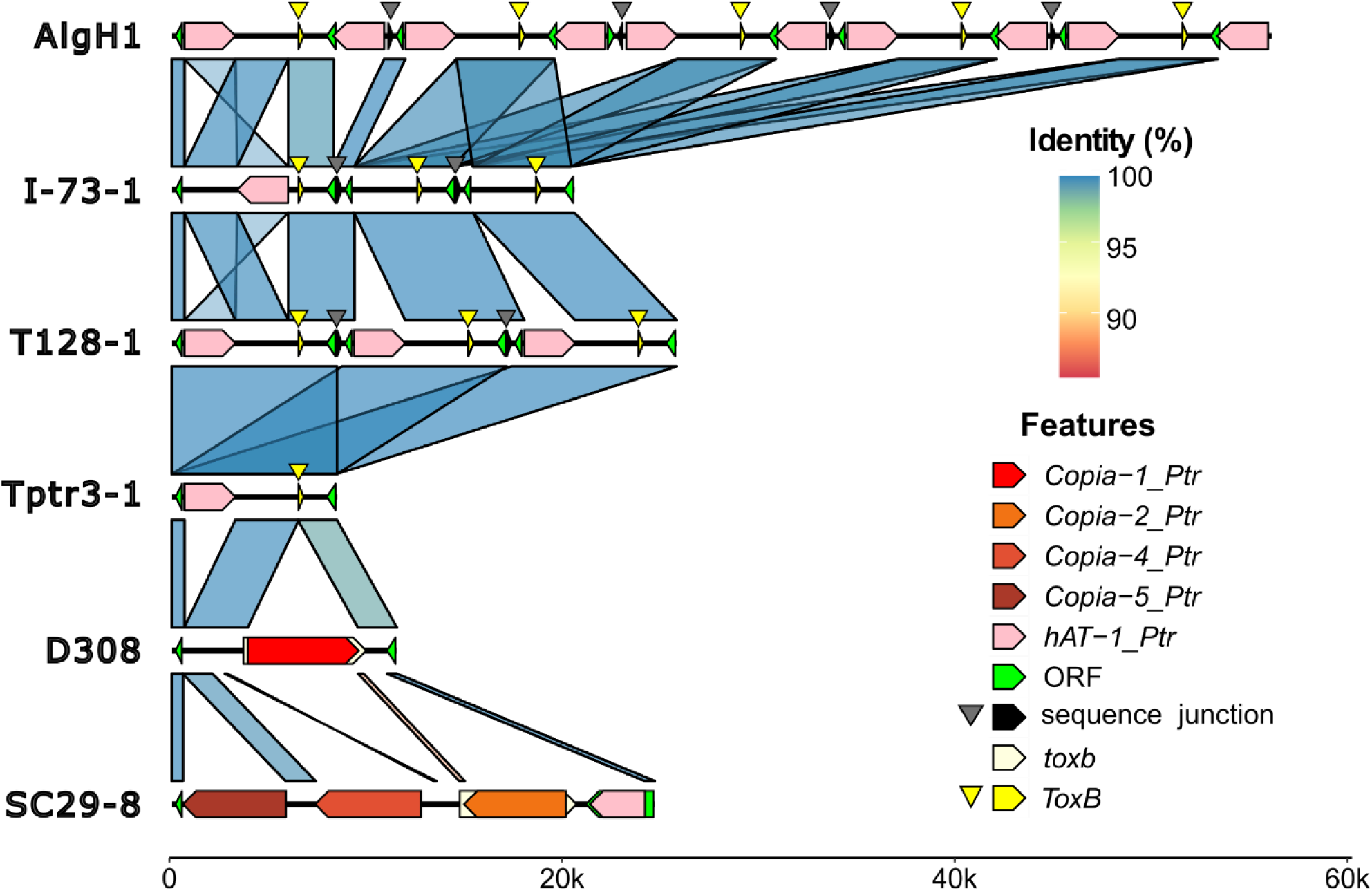
Linear alignments of the *ToxB/toxb* regions of Chr04 in *Pyrenophora tritici-repentis*. Colored arrows along the chromosome (black line) show *ToxB* (yellow), *toxb* (white), ORFs (green), transposons (reds and oranges), and the sequence junction between duplicated regions (black). Due to the alignment scale, vertical yellow and black arrows have been added to aid in visualizing the positions of *ToxB* and the sequence junction respectively. The linking blocks drawn between isolates show aligned DNA identity. Alignments less than 500 bp were omitted for clarity, as were most alignments between copies of the *hAT-1_Ptr* transposon, all of which are identical to each other. The position and number of *hAT-1_Ptr* transposons in the multi-copy isolates T128-1, I-73-1, and AlgH1 combined with the conserved edges suggest independent duplication events via the same mechanism. *ToxB* disruption via Copia retroelements is also visible in isolates D308 (*Copia-1_Ptr*) and SC29-8 (*Copia-2_Ptr*). An unedited version of the alignment can be found in Fig S5.

In summary, we obtained high confidence assemblies for four *ToxB* isolates where we could validate the *de novo* assembly with single-molecule long reads. Tptr3-1 carries a single *ToxB* copy, three isolates (I-34-1, I-73-1, and T128-1) carry a tandem triplication of *ToxB*. For AlgH1 alignment of the long reads to the *de novo* assembly supports five tandem *ToxB* copies, though these five tandem copies were not found in a single long-read. Isolates Alg3-24 and 92-171-R5 carry multiple copies of *ToxB* but not in the tandem nature observed in the other isolates. In these two isolates alignment of the long-reads to the *de novo* assembly showed that read coverage and contiguity near some *ToxB/toxb* sites was variable with coverage dropping to 10x in some regions.

While BLASTN results indicated that each *ToxB* copy was identical in the tandem duplications and the copies were closely spaced along the chromosome, they did not clarify whether the DNA in between the copies was also included in the duplication event. Leveraging the variation observed in copy number (i.e. isolates with one, three, or five tandem copies), the genomic region carrying *ToxB/toxb* was extracted and aligned to all other isolates, including self-alignments. These alignments revealed distinct 5’ and 3’ edges that defined the boundaries of the replicated sequence that included the *ToxB* gene. This sequence is referred to hereafter as the “*ToxB-*unit” (Fig. S4). An alignment of the *ToxB-*unit in isolates with five, three, and one copy plus two isolates with *toxb* is shown in Fig. 4. These alignments show that multi-copy isolates, carry perfect repeats of the *ToxB-*unit, but each perfect *ToxB-*unit repeat was separated by a conserved 251 bp “sequence junction” that was absent in any isolate with a single copy of *ToxB* or the inactive *toxb* (Fig. 4). The *ToxB-*unit includes two small open reading frames (ORFs), as well as a conserved hAT transposon that is 100% identical at the nucleotide level between different isolates (Fig. 4). In the triple copy isolates, I-73-1 and T128-1, the location of the hAT transposon varies which leads to repeats of the *ToxB-*unit that vary in size. In Tptr3-1, the only isolate carrying a single copy of *ToxB,* the unit was 8,416 bp from end to end. The smallest *ToxB-unit* is 5,798 bp in I-73-1 and the largest is in 11,434 in isolate AlgH1 as it contains two copies of the hAT transposon within the *ToxB*-unit. The ORF near the 5’ end of the *ToxB-*unit shared homology to protein domains in DUF3505 and C2H2-type zinc-fingers, while the second ORF near the 3’ shares homology with reverse transcriptase (RVT_1; PF00078) and RNases (RNase_H; PF00075). Both ORFs are quite small, 333 and 444 bp respectively, suggesting they may be gene fragments rather than fully functional genes. The relevance of each feature within the *ToxB-*unit is discussed in more detail below.

### *ToxB* is present in a dynamic landscape of repetitive elements

In annotating the features surrounding *ToxB* and performing detailed pairwise alignments between all isolates (not just those with tandem repeats shown in Fig. 4) it was clear that there were many unique sequence differences between different isolates with variable copies of *ToxB/toxb*. However, these differences were primarily driven by the nesting of other transposons near, or even within, the *ToxB* open-reading frame (Fig. 5; Table S1). After extensive manual curation of each sequence, we have classified 14 unique transposon insertions (5 DNA and 9 retrotransposons) near *ToxB* with an additional two insertions which appear to be transposons that we could not confidently classify (likely to be retrotransposons) (Table S1). We also identified three fragmented LTRs, two of which match the LTRs of the classified retrotransposons. Numerous copies of each were found throughout the *Ptr* genomes (Table S2). In order to assess whether the region carrying *ToxB* in Chr04 was richer in TEs compared to the rest of the genome we used two independent TE annotation software to identify TEs genome wide and then quantified what the TE density was in 50 kbp windows (Fig. S6). These annotations include simple repeats, low complexity regions, as well as transposons. Using I-73-1 as an example, the average number of annotations per 50 kbp bin across the whole genome is 15 for EDTA and 18 for EarlGrey. The average across the accessory region in Chr04 containing *ToxB* is 51 for EDTA and 35 EarlGrey indicating this region is repeat and transposon rich relative to the genome as a whole.

**Fig. 5.**
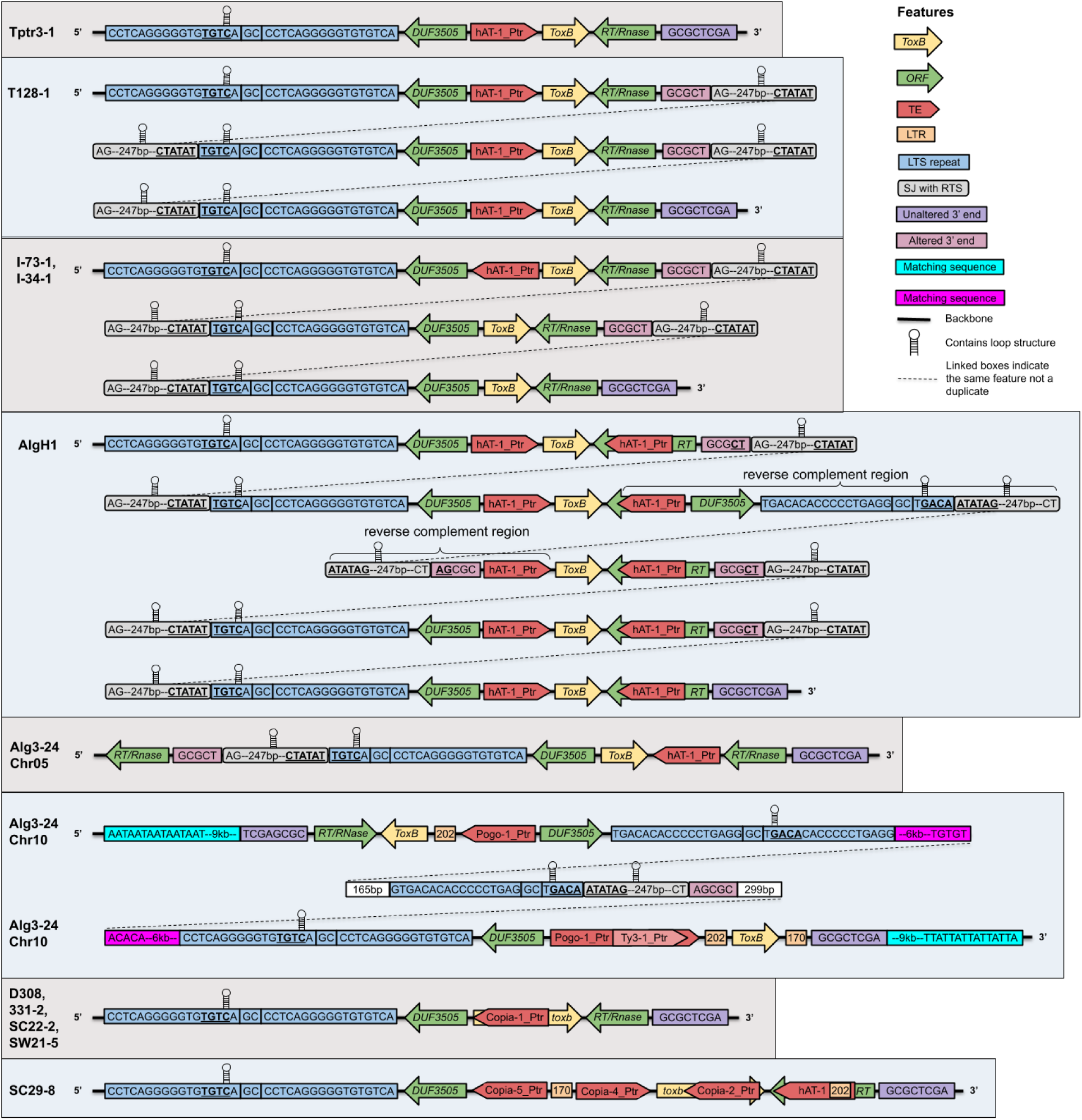
Schematic showing the diversity of eight *ToxB* haplotypes found in the isolates listed by name at the far left. Each individual or group of isolates with similar haplotypes are shown in a rectangular grey or blue box which alternate colours to improve visibility. Each repeated copy of the *ToxB-*unit is shown on one row, so boxes that contain three rows contain three tandem copies of *ToxB.* Overlapping sequences from each repeat are shown by the dashed line. Complete transposons, genes and other important sequences are shown in colored boxes or arrows according to the legend. Features that may be involved in the replication of the *ToxB-*unit are noted in the legend and are as follows: blue = 5’ left-terminal-sequence repeat; purple = original 3’ end; light pink = 3’ end in duplicated regions; grey = 251 bp sequence junction containing the right-terminal-sequence; green = open reading frames; red = transposons; orange = abandoned LTRs; loop icon = DNA hairpin structure is present in the region; dashed line = connected blocks are the same sequence. The sequence junction (SJ) is only present in the duplicated copies. Figure is not to scale and only reflects orientation and relative positions.

### Independent Copia transposon insertions create inactive toxb haplotypes

Recently, we showed that a single insertion of a 5.6 kbp Copia-like element, named *Copia-1_Ptr,* disrupts the 5’ end of the *ToxB* open-reading frame in isolate D308 (Hafez et al., 2024). To expand on this finding an additional nine *toxb* isolates were sequenced and aligned against the *ToxB* isolate Tptr3-1. Two *toxb* isolates, SW21-5 and SC22-2, contained the same *Copia-1_Ptr* insertion as D308 (Fig. 4 and 5) (Hafez et al., 2024). Previous Illumina sequencing for isolate SC29-1 demonstrated that the 3’ of *toxb* had many mutations compared to other isolates, which suggested that it may carry a different insertion (Gourlie et al., 2022; Hafez et al., 2024). The long-read assembly generated in this study confirmed this, as there is a novel 5.1 kbp insertion that disrupts the *ToxB* open-reading frame (Fig. 6). Within this 5.1 kbp insertion there are three large ORFs which encode a capsid protein (GAG), an integrase (INT) and retrovirus-related polyprotein (Pol), and reverse transcriptase (RT) domains. At the edges of the insertion, a four bp target site duplication was identified (5’-AAGT-3’) (Fig. 6). Adjacent to both TSDs were 165 bp LTRs. A search of RepBase revealed relatively high homology to the retrotransposon *Copia-3_PaNo-I* reported in *Parastagonospora nodorum* (67% identity over 46% of query) (Kojima, 2023). A pairwise alignment of this region with *Copia-1_Ptr* (which has disrupted the *ToxB* ORF in D308) was performed to determined similarity with the previously reported retrotransposon. Based on the alignment of the two retrotransposons and the large difference in LTR sequences the two Copia elements are unrelated (51% pairwise identity) (Fig. S7; alignment available on github) and following RepBase convention this transposon was named *Copia-2_Ptr*. *Copia-2_Ptr* is inserted at base 222 within the *ToxB* ORF, which places it further towards the 3’ end of the gene relative to the *Copia-1_Ptr* insertion in isolate D308 which was inserted eleven bases from the start of the gene. A second isolate, SC29-8, has an identical copy of *Copia-2_Ptr* inserted in the same location. These data indicate that the inactive *toxb* haplotype has arisen independently by the insertion of two different *Copia* transposable elements.

**Fig. 6.**
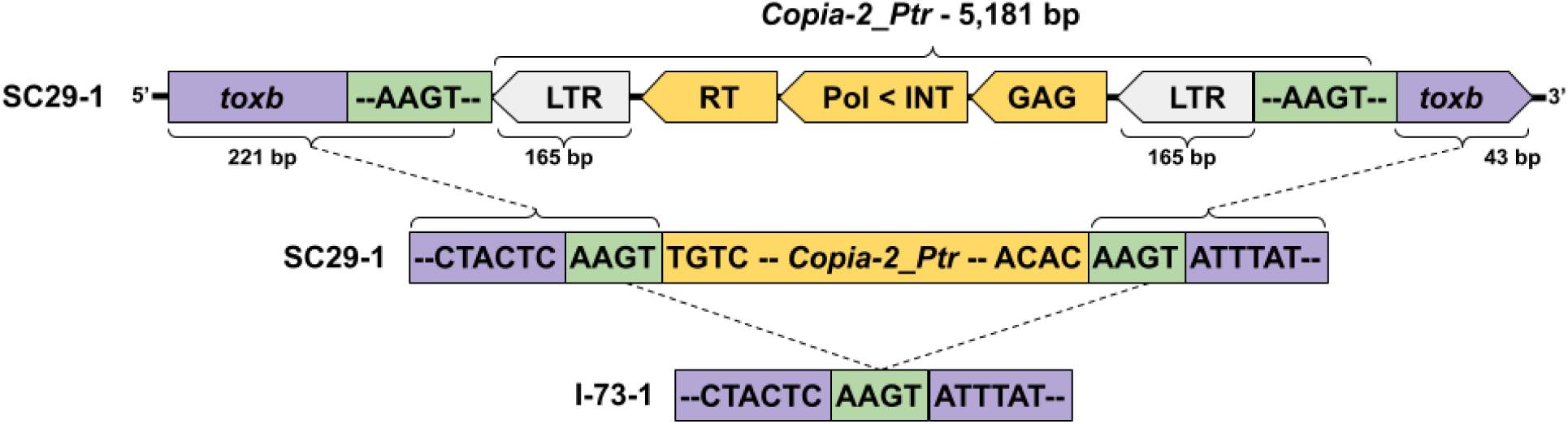
Schematic of a Copia-like transposon, *Copia-2_Ptr*, inserted within the 3’ end of the *ToxB* (purple) open-reading in isolate SC29-1 creating the inactive *toxb13* haplotype. A target site duplication (green) flanking long-terminal repeats (white) mark the transposon edges when compared with the non-transposon containing sequence of *ToxB* from isolate I-73-1 (purple). Open-reading frames within the transposon are marked by yellow arrows which show orientation (RT = reverse transcriptase; Pol = polyprotein; INT = integrase; GAG = capsid protein). Figure is not to scale.

### hAT transposon confirms multiple independent ToxB duplication events

The most commonly observed transposon insertion within the *ToxB*-unit is a 2,603 bp hAT transposon *hAT-1_Ptr* (Table S1; PV975802). This transposon is near *ToxB* in the majority of isolates, including all the multi-copy isolates and four of the single-copy (*ToxB/toxb*) isolates (Fig. 4 and 5). At the edges of the insertions are 6 bp TSDs (GCGGAC) that flank 17 bp TIRs. Four ORFs were identified within these TIRs and each ORF contained a conserved protein domain (Fig. S8). Copy numbers of this TE within entire genomes ranged from 0 to 35 copies with a stringent search criteria (90% ID over 90% query). A significantly larger element (5.6 kbp) was previously described by Manning et al., (2013), however upon inspection revealed that this larger transposon was *hAT-1_Ptr* with a second hAT transposon nested within it (3,278 bp; 6 bp TSD [5’-ATAGTC-3’]).

The varied locations of *hAT-1_Ptr* offers key insights into the history of the *ToxB-*unit, in particular when comparing isolates with the highest quality assemblies; the single copy Tptr3-1, triple copies T128-1, I-73-1 and I-34-1, and quintuple copy AlgH1. The single *ToxB* isolate Tptr3-1 and the triple-copy isolate T128-1 both contain *hAT-1_Ptr* in the exact same relative position within the *ToxB-*unit, indicating that the T128-1 is a perfect triplication of the sequence observed in Tptr3-1 (Fig. 4 and 5). However, in isolates I-73-1 and I-34-1, only a single copy of *hAT-1_Ptr* is present and it is located in only one of the three *ToxB-*units. This indicates that in these isolates the triplication of *ToxB* must have occurred prior to the insertion of *hAT-1_Ptr*. Finally, in AlgH1, which has five *ToxB*-units, there are two copies of *hAT-1_Ptr* present (Fig. 4 and 5). Each *ToxB*-unit contains the two copies of *hAT-1_Ptr* at the same position in each unit, which indicates that *hAT-1_Ptr* inserted prior to the amplification of the *ToxB-*unit. The different conserved locations of *hAT-1_Ptr* in the different isolates is clear evidence that tandem duplication of *ToxB-*unit has occurred multiple independent times in *Ptr* (a minimum of three independent events in the isolates sequenced thus far). Each independent amplification event was flanked by conserved edges of the *ToxB-*unit, which suggested that there was a common mechanism driving the independent amplifications of this region.

Given the prevalence of transposons in the region, including *hAT-1_Ptr,* we first explored whether there was any evidence that any of these transposons drove the amplification of the *ToxB*-unit. The edges of the *ToxB*-unit contained completely unmatching sequences, meaning they were neither long-terminal repeats (LTR) nor terminal inverted repeats (TIR) common to most transposon families. Additionally, no discernable target-site duplication (TSD) were identified when compared to isolates without *ToxB/toxb*. This excluded the *hAT* transposase located within the duplicated sequence as the driver of the duplication, as well as most described Class I or Class II transposons. In between each tandem repeat of the *ToxB* region, there is a small sequence of 251 bp which is referred to as the sequence junction (SJ) (Fig. 5 and S4). The SJ is only present in isolates with multiple tandem copies of the *ToxB-*unit and absent in single copy *ToxB/toxb* isolates and therefore seems to be important for the accumulation of tandem arrays of the *ToxB-*unit (Fig. 5). The SJ is also highly abundant (28 to 85 copies) in all *Ptr* genomes with >98% nucleotide similarity over 90% of the query (Table S1; S2). Due to the prevalence of this sequence throughout the genome, we selected random instances for closer inspection. In most cases, the SJ appears to be a conserved domain located within reverse transcriptases or LINE transposable elements (a type of retrotransposon). However, there are instances where the SJ were not part of any ORF, annotated gene, or transposon. BLAST searches with the SJ nucleotide sequence on NCBI and UniProt yielded limited similarity to reverse transcriptase, RNaseH1, hAT-Tnp-dimer, and DUF3824 sequences (∼40 - 60% identity). Intersection between the SJ location and transposons annotated with EDTA or EarlGrey resulted in many LINE element overlaps. How each of these elements are phylogenetically related to the SJ associated with the duplication of *ToxB* is shown in Figure S9. Based on these findings it is likely the SJ is a conserved domain within some retrotransposons and the truncated version found within the *ToxB* duplicated regions plays a critical role.

Other genomic mechanisms of duplication were also considered, such as microhomology break induced replication and breakage fusion bridge cycles, however, given the restricted size of the duplication and the different number of tandem arrays of the *ToxB-*unit micro-homology repair mechanisms did not seem to fit with the observed sequence data. We therefore expanded our search for a mechanism to other less common transposon families.

### Diagnostic features support a Helitron-like element associated with *ToxB*

Helitrons are DNA transposons that are hypothesized to use a rolling-circle mode of replication. These transposons lack TSDs, are structurally asymmetrical (i.e. non-matching edges), and can produce copy-paste style tandem (or near-tandem) duplications (Kaptinov and Jurka, 2001; 2007; Thomas and Pritham, 2015). Most Helitrons are non-autonomous, utilizing existing molecular machinery to mobilize themselves, however some do carry Replication initiator/helicase (RepHel) and/or replication protein A (RPA) domains (Boa and Jurka, 2013; Thomas and Pritham, 2015; Grabundzija, et al., 2016). Furthermore, Helitrons are known to capture genes and/or gene fragments, which fits with the duplication of *ToxB* and other observed ORFs within the replicon. To initiate replication, canonical Helitrons contain a 5’ ‘TC’ dinucleotide, where the Rep/Hel protein binds, known as the left-terminal sequence (LTS). The Rep/Hel protein replicates the DNA sequence from this 5’ LTS until it reaches a 20-40 bp hairpin loop followed by ‘CTRR’ (R = A or G), known together as the right-terminal sequence (RTS) (Kaptinov and Jurka, 2007; Thomas and Pritham, 2015; Grabundzija, et al., 2016). Helitrons within the fungal kingdom are very poorly characterised, though there is one study with *Fusarium oxysporum* that defines several novel Helitron families. These Helitrons, have two hairpin structures both at the 5’ LTS and 3’ RTS. Moreover, the starting 5’ initiating sequence can vary, with two di-nucleotides (TC and TA) as well as a tri-nucleotide (TGC) (Chellapan et al., 2016). The *F. oxysporum* RTS also differ from other described termination sequences and two different five base pair termination sequences have been described. These general features of canonical versus *Fusarium* specific Helitrons are summarised in Fig. S10.

In *Ptr* the 5’ end of the *ToxB-*unit in both single and multi-copy isolates contains two 17 bp repeats separated by a ‘GC’ dinucleotide (i.e. 36 bp total) (Fig. 5). This 36 bp contains several potential ‘TC’ dinucleotide initiation sites. However, in isolates with tandemly duplicated copies, this region has been truncated to 24 bp and consistently starts with ‘TGTC’ (Fig. 5). This was observed in multiple different *ToxB* haplotypes, which as described above are likely the result of independent amplification events of the *ToxB-*unit. The 3’ end of the *ToxB*-unit on internal copies was ‘CTATAT’ (i.e. ‘CTRTAT’). This partially matches the canonical 3’ CTRR termination sequence. As described, the canonical RTS of Helitrons contains a ∼20 to 40 bp GC dominant stem loop which begin 30 to 60 bp upstream of the 3’ ‘CTRR’ sequence. The *ToxB-*unit RTS sequence was analyzed using VectorBuilder to establish whether sequences upstream of the 3’ end formed a stem loop structure (Fig. S11). One stem loop was found 72 bp upstream of the 3’ ‘CTATAT’ sequence. This loop has a ΔG of -5.90 and is GC dominant. A ΔG value of -5.90 indicates loop formation is spontaneous, energetically favorable and moderately stable. Interestingly, the strength of this loop exactly matches that of *FoHeli1* in *F. oxysporum* which suggests this loop strength is sufficient to enable replication (Fig. S11) (Chellapan et al., 2016). The 5’ end also contained a stem loop with a ΔG of -6.20, again matching the description of fungal Helitrons described in *Fusarium* (Fig. S11).

Some Helitrons contain RepHel and/or RPA domains which facilitate their mobility, however we did not observe any within the putative replicon edges. Functional gene annotations revealed at least ∼80 Helicase genes are present within the *Ptr* genome, with at least three associated with Helitrons (i.e. Pif1-like helicases) (Heringer and Kuhn, 2021; Castanera et al., 2014; Thomas et al., 2014). Several DNA2-NAM7 helicases were also identified which have been linked to rolling-circle replication in viral plant pathogens but not Helitron activity specifically (Sahu et al., 2010; Maredza et al., 2016). Most annotated helicases were of the DEADbox class which have limited support for Helitron involvement (Tuteja, 2003). Gene annotations also identified at least two RPA-like genes. Based on the presented features and the confirmation that replication has occurred independently multiple times we conclude that a **<u>H</u>**elitron-**<u>L</u>**ike **<u>E</u>**lement, which we’ve named *ToxB-HLE*, is a likely candidate to explain duplication of *ToxB* in the genome of *P. tritici-repentis*. A potential model of replication is discussed below (Fig. 7) .

**Fig. 7.**
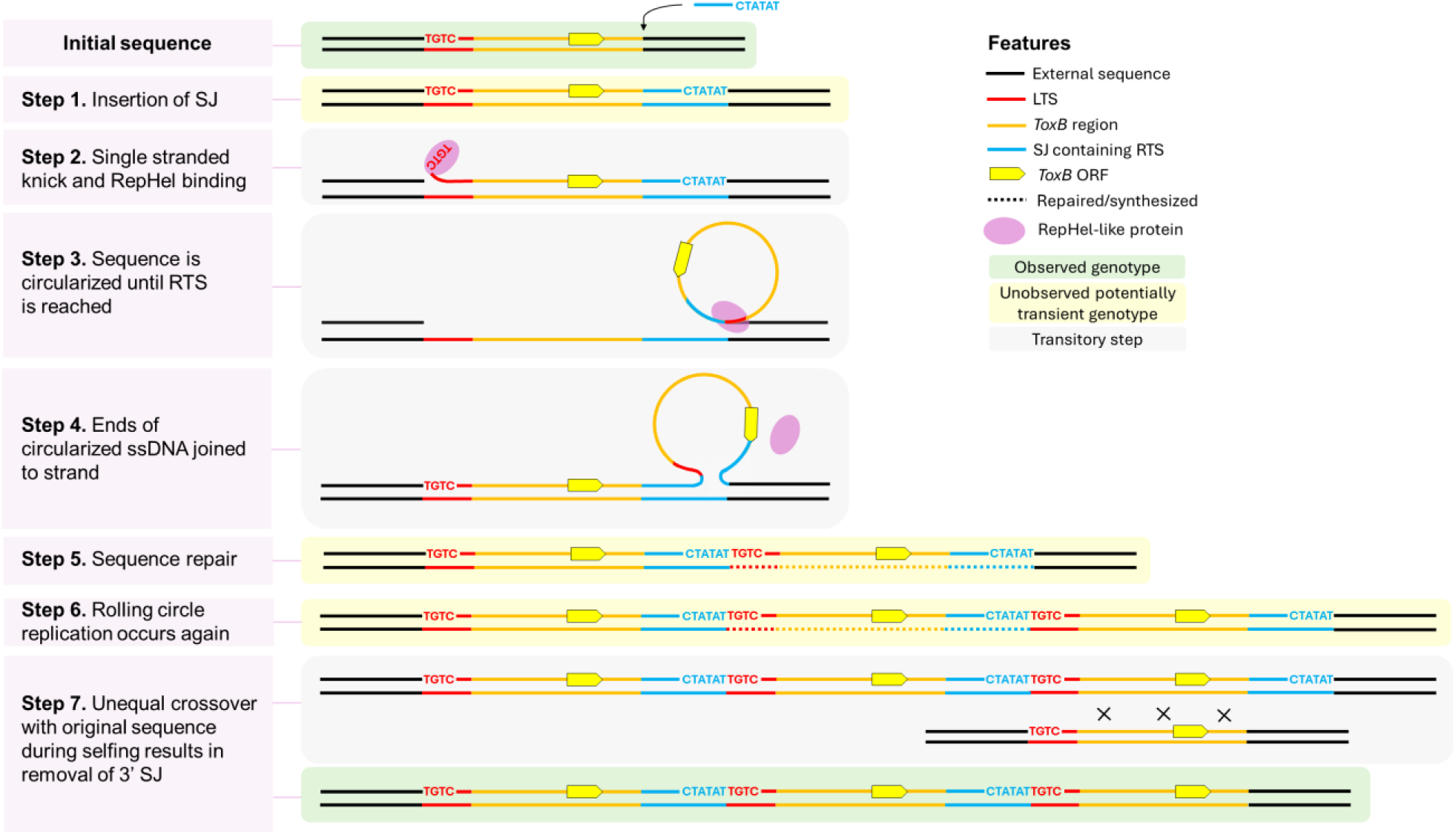
Steps which could lead to the observed genotypes through insertion, rolling-circle replication, and unequal crossing over. In this model, *ToxB-HLE* is composed of the *ToxB* region (orange) containing the left-terminal sequence ‘TGTC’ (red), and the sequence junction (blue) which contains the right-terminal sequence ‘CTATAT’ as well as the requisite loop structure (not pictured here; see Fig. S11). Observed genotypes are on a green background. Figure is partially adapted from Thomas and Pritham (2015), Chellepan et al., (2016), and Grabundzija et al., (2016).

## Discussion

The evolution of effectors in plant pathogens is a critical component to maintaining their survival and fitness. Therefore, understanding which molecular mechanisms drive adaptive changes in these genes is a key part of developing long-term disease management strategies. In the tan spot pathosystem, past works have identified two major necrotrophic effectors which have been characterized and studied for decades due to their economic impacts (ToxA and ToxB) (reviewed in Aboukhaddour 2023; Hafez et al., 2024). For over twenty years, the mechanisms driving the amplification of *ToxB* has remained largely unexplored (Strelkov et al. 1999; Martinez et al., 2001; Lamari et al., 2003). It is established that isolates with a high-copy number of *ToxB* have been shown to be more virulent on wheat (Strelkov et al., 2002; Martinez et al., 2004; Aboukhaddour et al., 2012). While increased copy number of other effectors, such as NIP1 in *Rhyncosporium commune,* have also been linked to increases in virulence, it is rare to identify a putative mechanism that drives gene amplification within a genome (Mohd-Assaad et al., 2019). Here we provide a comprehensive examination of the evolution of the *ToxB* locus, providing evidence that a non-autonomous Helitron-like-element, *ToxB-HLE*, is responsible for multiple independent duplication events of this effector. At the same locus, we also show that *ToxB* has been inactivated in at least two independent events through the insertion of Copia retrotransposons, highlighting the dual nature that transposons play in the evolution of virulence. This work is an important example, in a growing number of studies, of how recent transposon activity appears to drive local adaptation in different fungal populations within the same species (Wang et al., 2021; Feurtey et al., 2023).

### A place for change - transposons create a volatile landscape near *ToxB*

Among 17 *ToxB/toxb* isolates investigated for this study, this effector was located ∼400 to 600 kbp from the end of Chr04 in 12 isolates. Earlier long-read sequencing of *Ptr* using two isolates (I-73-1 and D308) suggested this region may be an ancient and derelict *Starship* (dubbed *Icarus*) due to the presence of several known *Starship* cargo genes (Gourlie et al., 2022). This previous finding is supported by the extended sequencing performed for this study where this region is completely absent in all but one non-*ToxB/toxb* isolates. Large accessory regions (100-200 kb) are one of the key signatures used to identify putative *Starships* (Gluck-Thaler and Vogan, 2024). In previous work, several *Starship* accessory genes were identified in this accessory region along with a number of TEs from other various transposon classes (Gourlie et al., 2022). This region however appeared to lack the signature tyrosine recombinase transposase that would provide more definitive evidence that this region was a *Starship*. Further investigation of this hypothesis, and the origins of the *ToxB* locus in *Ptr*, is an exciting area for future work, especially in the context of the recent discovery on the prevalence and diversity of *Starship* transposons within the *Pezizomycotina* (Gluck-Thaler et al., 2022; Gluck-Thaler and Vogan, 2024; Urquhart et al., 2024). *ToxB/toxb* is unique when compared to other fungal effectors, as homologs of this effector are also found in other closely related species; for example the bromegrass pathogen *Pyrenophora bromi* (Andrie and Ciuffetti, 2011; Hafez et al., 2024). Further sequencing of this related pathogen could provide further insight into the evolutionary history of this accessory region carrying *ToxB/toxb* and its presence or not in a larger *Starship* TE.

Accessory genomic regions in fungal pathogens are known to be enriched in virulence genes and contain many repetitive elements, which is hypothesised to lead to higher rates of genomic rearrangement and mutations (Wöstemeyer and Kreibich, 2002; Faino et al., 2016; Bertazzoni et al., 2018). This general observation led to the development of the “two-speed” or “multi-speed” genome hypotheses for fungal genomes, that these repeat-rich regions were hotbeds for the accumulation of mutations that could then be selected for under environmental or host stresses (Raffaele et al., 2010; Croll and McDonald, 2012; Dong et al., 2015; Torres et al., 2020). This hypothesis has been recently refined to refer to genome compartments, rather than speeds, as it remains unclear whether genome regions with a high density of transposons do indeed undergo more rapid change, or instead, that these regions have higher levels of observed rearrangement/mutations due to relaxed selection (Torres et al., 2020). An abundance of transposons were found very near to *ToxB* creating a landscape of repetitive elements that has led to several structural rearrangements within this region. These rearrangements are independent of the observed *ToxB* tandem duplication events, and in some isolates were so complex that even with long-HiFi PacBio reads they could not be fully resolved in the genome assemblies. For example, in isolates Alg3-24 and 92-171R5, it is possible that these repeats led to the translocation of *ToxB* to other chromosomes. In *Ptr,* transposon activity could be potentially exacerbated by the fact *Ptr* appears to lack RIP, an important defense mechanism in fungi which can deactivate transposons (Gladyshev, 2017; Gourlie et al., 2022). Overall, the observations of substantial rearrangements and TE density in this region in multiple long-read assemblies, supports the long-standing view that fungi are largely tolerant of extensive chromosomal rearrangements.

While tolerant of genome instability, there is clear evidence that TEs do not only promote adaptive genomic changes, as we also observe multiple independent pseudogenizations of *ToxB* through the insertion of two different Copia retrotransposons. Effector gene deletion or pseudogenization is more commonly reported for genes with avirulence functions, whereby recognition leads to plant immunity (Fouché et al., 2018). Virulence gene disruption by transposons have been reported in other fungal species such as *Fusarium oxysporum*, where the Avr gene *SIX1-H* was shown to be deactivated by a hAT transposon (Rep et al., 2005). In this case, the disruption is adaptive as it prevents recognition of the pathogen by the plant immune system. However, for necrotrophic effectors, like *ToxB*, pseudogenization leads to a loss of virulence (Powell et al., 2008; Inami et al., 2012).

### Structural features support Helitron mediated *ToxB* duplication

There are a diverse set of described mechanisms that can drive gene duplication, some of which are difficult to tease apart ex post facto (e.g. TE-mediated gene capture, template-switching, break-fusion-bridge, illegitimate cross overs, etc.) (Reams and Roth, 2015). Using careful pair-wise alignments between different individuals and different copies of the *ToxB* repeat, we have identified a number of structural features that provide strong evidence for a controlled and repeatable replication of a defined sequence. We propose that this is made possible by a novel, non-autonomous Helitron named *ToxB-HLE*.

Relative to other transposable elements, Helitrons are poorly understood and difficult to identify (Kaptinov and Jurka, 2001; 2007; Thomas and Pritham, 2015; Grabundzija et al., 2016; Chellapan et al., 2016). As such, they are likely to be underrepresented in transposon annotation pipelines and databases, particularly if they do not conform to the described Helitron model (Xiong et al., 2014). The identification of large families of Helitrons in plants has been facilitated by semi-conserved sequences at the termini of these elements, and most bioinformatic software trained to annotate Helitrons relies on these conserved sequences (Ou et al., 2019; Baril et al., 2024). However in fungi, no such conserved sequences have been identified, making Helitron identification much more difficult (Chellapan et al., 2016). Identifying Helitrons within fungal genomes could be important as they have consistently been associated with the generation of novel gene sequences that are actively transcribed (Kapitonov and Jurka 2001; 2007; Sweredoski et al., 2008; Fu et al., 2013). In the oyster mushroom *Pleurotus ostreatus*, HELPO1 and HELPO2 Helitron families were identified capturing genes with unknown functions (Castanera et al., 2014). HELPO1 also experienced incursion by retrotransposons within the Helitron boundaries, creating chimeric transposons similar to what we have observed within *Ptr*’s *ToxB-HLE* (Castanera et al., 2014). Similarly, a genome-wide study of *Rhynchosporium commune*, found a small number of Helitrons associated with an increased copy-number of captured genes, and in *Fusarium*, 27 gene captures by Helitrons are reported to have resulted in increased copy-numbers (Chellapan et al., 2016; Stalder et al., 2022). Also in *F. oxysporum*, Helitrons were reported to be very near to pathogenicity-related genes (e.g. SIX9a, SIX9b, SIX6, ORX1-like, and ARG1) though none of these were captured (Chellapan et al., 2016). Other researchers have suggested that ectopic recombination or unequal crossing over caused by the presence of Helitrons may have played a role in the divergence of pathogenic races and the creation of accessory elements in *F. oxysporum* (Schmidt et al., 2013; Vlaardingerbroek et al., 2016; Biju et al., 2017). Together these examples suggest that Helitrons are prevalent in fungal genomes and playing important roles in gene copy number variation. Therefore, describing and classifying novel fungal Helitron features, and ideally integrating these into available TE detection software, would greatly enhance our ability to comprehensively describe the frequency of Helitron-mediated gene capture.

The first notable aspect of *ToxB* replication is the tandem nature of the duplication. Helitrons transpose via rolling-circle replication which is capable of generating such tandem repeats (Thomas and Pritham, 2015; Grabundzija et al., 2016; Xiong et al., 2016; Dias et al., 2016). While Helitrons do not possess repetitive edges as often seen in other transposons (i.e. LTRs in retrotransposon or TIRs in DNA transposons) they do possess conserved sequences at their 5’ and 3’ ends as well as diagnostic step loops which are important for their replication. In *F. oxysporum,* Helitrons have been reported to be different from canonical elements described in other kingdoms (Chellapan et al., 2016). Standard Helitrons contain a ‘TC’ dinucleotide at their 5’ end, while the *ToxB-HLE* 5’ end contains a conserved ‘TGTC’ sequence. This 5’ end more closely matches Helitrons in *F. oxysporum* where a 5’ ‘TA’ or ‘TGCCT’ start sequence have both been described (Chellapan et al., 2016). For the 3’ right terminal sequence, canonical Helitrons have a conserved 20-40 bp hairpin loop followed by ‘CTRR’ (R = A or G), which again differs for what has been described for fungi. *ToxB-HLE* does have an energetically favorable 3’ stem loop occurring 72 bp upstream of the putative 3’ termination sequence ‘CTATAT’. This putative end for *ToxB-HLE* again more closely matched the described 3’ Helitron end for *F. oxysporum* (‘CTCCTGT’) (Chellapan et al., 2016). In addition to the 3’ stem loop, *ToxB-HLE* also contains a 5’ stem loop (Thomas and Pritham, 2015). The presence of 5’ stem loop could indicate *ToxB-HLE* belongs to the subgroup Helitron2 (or possibly Helentrons), however asymmetrical inverted repeats (ATIR) associated with Helitron2 were not found (Boa and Jurka, 2013).

Insertion site preference is another common way transposons can be classified into specific families. For example, Helitron1 elements have shown a bias for insertion between ‘A/T’ dinucleotides, while Helitron2 and Helentrons are more commonly observed at ‘T/T’ or ‘T/C’ insertions (Thomas et al., 2014; Thomas and Pritham, 2015; Grabundzija, et al., 2016; Chellapan et al., 2016). However, this does not appear to be a hard rule, as Grabundzija et al., (2016) found *Helraiser* insertions between almost all possible dinucleotide pairs. In the case of *ToxB-HLE,* insertions appear to occur between ‘T/T’ dinucleotides (Fig 5). The lack of the RepHel/RPA domains in *ToxB-HLE* is notable but it has been established that the vast majority of Helitrons are non-autonomous and therefore most will lack these domains (Boa and Jurka, 2013). We were able to identify helicases and RPA genes within the *Ptr* genome which may support *ToxB-HLE* replication. It bears repeating here that in the majority of examined isolates, the duplication of *ToxB* has occurred in a conserved region, and most importantly, the different numbers and orientations of the *hAT-1_Ptr* transposon point to multiple independent replication events through a common mechanism.

### A model for the amplification of *ToxB*

Combining all information together, we’ve provided a potential model which outlines the steps which may have led to the observed genotypes (Fig. 7). In this model, the initial sequence is observed in a single copy *ToxB* containing genome (represented by isolate Tptr3-1). An insertion of the SJ then occurs in Step 1 which brings with it the requisite RTS and stem loop. A single copy of *ToxB* with the SJ is, so far, an unobserved genotype that may be transient in nature. Based on previous works, Steps 2 through 6 broadly outline how Helitrons are thought to undergo rolling-circle replication and create tandem duplications (Thomas and Pritham, 2015; Chellapan et al., 2016; Grabundzija et al., 2016; Dias et al., 2016; Harmer et al., 2022). Rolling circle replication is followed by an unequal crossover event which removes the terminal SJ and may continue to amplify *ToxB* copy numbers. Tandemly duplicated regions become more likely to undergo unequal crossing events due to the increased homology (Anderson and Roth, 1977; Silver, 2001; Taylor and Raes, 2004). These events are made more likely through the fact that *Ptr* is homothallic, and can cross with itself, providing ample opportunity for such unequal crossings to occur. Interestingly, it has also been suggested that homothallics may themselves arise from the unequal crossing over of heterothallics (Yun et al., 1999). For example, in some *Neurospora* species, unequal crossing over of mating type genes facilitated by LTR retrotransposons appear to have prompted the transition from heterothallism to homothallism and this may be the cause of *Ptr*’s homothallism as well (Gioti et al., 2012). There is no hard limit on the number of copies which could be created through the activity of *ToxB-HLE* combined with unequal crossing over which explains isolates such as AlgH1 with 5 copies or other isolates with as many as 10 copies reported (Strelkov et al. 2005; Amaike et al. 2008; Aboukhaddour et al. 2012). This model is not functionally validated, nor is the activity of *ToxB-HLE*. During this investigation we considered microhomology mechanisms, other transposon classes, and other Helitron replication models. For example in IS91 found in bacteria, deletion of the RTS leads to the accumulation of tandemly duplicated IS91 copies as opposed to what is proposed here that is the insertion of the SJ that drives the tandem duplication (Mendiola et al., 1994; Dias et al., 2016).

The activity of *ToxB-HLE* may not be the only mechanism driving changes in *ToxB* copy-number. As the region expanded with higher numbers of nested transposons in some isolates, other unexplained mechanisms may also replicate *ToxB* as seen in Alg3-24 where features of *ToxB-HLE* are present but scattered within a much larger duplicated region on Chr10 and across many other small unassembled contigs (Fig. 5; Table 1). The structure surrounding copies in Alg3-24 could also be explained by genomic rearrangements which have broken up the previously tandem copies over time. This scenario seems likely, as more evidence builds showing strain heterogeneity and genomic rearrangements occur in isolates stored for long time periods as is the case with Alg3-24 (collected in 1993) (Keller, 2017; Lofgren et al., 2022; Gluck-Thaler et al., 2025). Additionally, in isolate 92-171R5 we observed two copies of *ToxB* but did not find any signatures of *ToxB-HLE* in the region, rather, the duplicated regions are flanked by undefined LTR retrotransposons (as annotated by EDTA) (Fig. S12). The relatively poor coverage of this region in 92-171R5 combined with the abundance of transposons makes it difficult to draw confident conclusions for this isolate. However if the assembly is accurate, its possible the duplication of *ToxB* was driven by retrotransposition in 92-171R5 rather than *ToxB-HLE*. Previous work found that 92-171R5 is evolutionarily divergent to other *Ptr* isolates, and so it is not surprising to see different mechanisms may be at play in the evolution of *ToxB* (Gourlie et al., 2022).

This work highlights the double-edged nature of transposable elements which can create gene duplications, cause gene disruptions and facilitate genomic rearrangements (Fouché et al., 2022). In fungal pathogens in particular, gene duplications have been reported in various metabolic systems, however, rarely is a specific of mechanism identified (Powell et al., 2008; Stalder et al., 2022). Here we have provided a large body of evidence to support the hypothesis that multiple types of transposons act to both amplify and disrupt an important necrotrophic effector in a global plant pathogen. In *Ptr*, it is unclear how long these duplicated versions of *ToxB* will be maintained before they are either lost, pseudogenized, or retained. Copy retention and conservation is likely linked to the increased fitness the extra *ToxB* copies may provide on susceptible wheat hosts. However, should additional copies no longer provide an advantage, such redundancy is unlikely to be preserved and additional copies would be free to mutate and diverge (Silver, 2001; Taylor and Raes, 2004). Therefore, it is possible that in the future, *ToxB* copies may evolve to become novel effectors with altered functions or new host target sites. As such, continued investigation and world-wide screening of *ToxB* and its homologs can provide a framework for understanding novel necrotrophic effector formation.

## Acknowledgments

We would like to acknowledge the late Dr. Lakhdar Lamari, whose pioneering work on *Pyrenophora tritici-repentis* laid the foundation for *ToxB* research. Several isolates used in this study were collected by him during his surveys across the Fertile Crescent, North Africa, and the Caucasus regions. We also would like to acknowledge Tara Rintoul (Canadian Collection of Fungal Cultures, AAFC, Canada (DAOMC)), Sana Kamel and Mejda Cherif (University of Carthage, Tunisia), Lise Nistrup Jørgensen (Aarhus University, Denmark), and Kaori Nakajima (Mie Prefectural Agricultural Research Institute, Matsusaka, Japan) for providing a number of the isolates in this study. We thank Mouldi Zid for his help in growing and maintaining the isolates as well as bioassays.

Funders: Agriculture and Agri-Food Canada, Alberta Grain, and Saskatchewan Wheat Development to RA. The funding bodies were not involved in the design of the experiments, data collection, analysis, or interpretation, or in the writing of this manuscript. MCM is supported by a UKRI Future Leaders Fellowship (MR/Y01717X/1)

## Data Availability

Genomic data was deposited in NCBI under BioProject PRJNA803191. Scripts and other files available on Github at https://github.com/fungal-spore/ToxB_replication.

## Supporting Information

**Fig. S1** LASTZ alignment showing variable size of accessory region compared to isolates which do not contain *ToxB/toxb*.

**Fig. S2** Dotplots showing single reads (y-axes) which cover all copies of *ToxB* in the assemblies (x-axes) of four multicopy isolates.

**Fig. S3** Read coverage around *ToxB* in isolate Alg3-24.

**Fig. S4** Example of self-alignments done with isolate T128-1 which were used to locate edges of the *ToxB*-unit.

**Fig. S5** Linear alignments between ToxB-units in representative isolates, unedited version of Figure 4.

**Fig. S6** Transposon density plots of I-73-1 as a representative example.

**Fig. S7** Dotplot alignment of *Copia-1_Ptr* and *Copia-2_Ptr*, disruptors of the *ToxB* ORF.

**Fig. S8** Schematic of *hAT-1_Ptr* showing target site duplications, terminal inverted repeats, and defining protein domains.

**Fig. S9** Phylogeny of transposable elements which contain the sequence junction.

**Fig. S10** Generalized schematics of known Helitrons and *ToxB-HLE*.

**Fig. S11** Loop structures found near the terminal edges of *ToxB-HLE*.

**Fig. S12** Geneious annotations surrounding copies of *ToxB* (green arrows) in isolate 92-171R5.

**Table S1** Details of transposons and insertions near or within the *ToxB* ORF in P*. tritici-repentis*.

**Table S2** Counts of transposons and the sequence junction throughout the genomes of *P. tritici-repentis*. Counts based on BLASTN 90% identity over 90% of the sequence.

**Methods S1** Details of PacBio sequencing methodology.

