## Supplemental Table S2 for "The virulence gene *ToxB* is both amplified and disrupted by transposons in the wheat pathogen *Pyrenophora tritici-repentis*"

Table S2. Counts of transposons and the sequence junction throughout the genomes of *P. tritici-repentis* . Counts based on BLASTN 90% identity over 90% of the sequence. Isolate names are followed by B, b, or N representing ToxB, toxb, or neither respectively

| Isolate | Transposon copy count | | | | | | | | | | | | | | |
|---|---|---|---|---|---|---|---|---|---|---|---|---|---|---|---|
| | ***Copia-1_Ptr*** | ***Copia-2_Ptr*** | ***Copia-3_Ptr*** | ***Copia-4_Ptr*** | ***Copia-5_Ptr*** | ***Copia-6_Ptr*** | ***Copia-7_Ptr*** | ***hAT-1_Ptr*** | ***hAT-2_Ptr*** | ***hAT-3_Ptr*** | ***hAT-4_Ptr*** | ***Pogo-1_Ptr*** | ***Ty3-1_Ptr*** | ***Ty3-2_Ptr*** | **SJ** |
| **107224_b** | 3 | 0 | 4 | 0 | 0 | 1 | 6 | 0 | 0 | 1 | 0 | 8 | 0 | 1 | 53 |
| **331-2_b** | 12 | 4 | 5 | 15 | 19 | 0 | 25 | 26 | 0 | 44 | 54 | 24 | 0 | 7 | 53 |
| **90-2_b** | 0 | 0 | 2 | 0 | 0 | 0 | 4 | 0 | 0 | 0 | 0 | 7 | 0 | 0 | 58 |
| **92-171-R5_** | 9 | 0 | 2 | 0 | 2 | 21 | 2 | 0 | 0 | 5 | 12 | 38 | 0 | 0 | 58 |
| **Alg3-24_B** | 1 | 5 | 0 | 19 | 9 | 1 | 24 | 20 | 1 | 21 | 34 | 28 | 6 | 6 | 63 |
| **AlgH1_B** | 10 | 2 | 5 | 18 | 13 | 0 | 24 | 33 | 0 | 58 | 62 | 26 | 0 | 8 | 48 |
| **D308*_b** | 9 | 3 | 5 | 14 | 15 | 0 | 25 | 16 | 0 | 45 | 63 | 23 | 0 | 6 | 30 |
| **Den17_b** | 3 | 2 | 1 | 21 | 9 | 0 | 28 | 20 | 0 | 26 | 48 | 25 | 1 | 5 | 44 |
| **G9-7_b** | 4 | 0 | 1 | 0 | 10 | 3 | 2 | 0 | 0 | 0 | 0 | 17 | 0 | 4 | 85 |
| **I-33-16_N** | 5 | 3 | 1 | 18 | 9 | 0 | 22 | 19 | 0 | 22 | 41 | 30 | 3 | 5 | 42 |
| **I-34-1_B** | 16 | 2 | 5 | 16 | 16 | 0 | 19 | 28 | 0 | 35 | 56 | 28 | 0 | 12 | 28 |
| **I-73-1_B** | 15 | 2 | 5 | 15 | 14 | 0 | 19 | 25 | 0 | 39 | 51 | 24 | 0 | 5 | 29 |
| **K11_N** | 0 | 0 | 0 | 0 | 0 | 0 | 0 | 0 | 0 | 1 | 31 | 14 | 1 | 11 | 51 |
| **K6_N** | 0 | 0 | 0 | 0 | 0 | 2 | 0 | 0 | 0 | 0 | 17 | 11 | 3 | 9 | 46 |
| **K9_N** | 0 | 0 | 0 | 0 | 0 | 3 | 0 | 0 | 0 | 1 | 30 | 13 | 2 | 10 | 47 |
| **SC22-2_b** | 12 | 4 | 5 | 14 | 19 | 0 | 25 | 23 | 0 | 42 | 53 | 26 | 0 | 7 | 64 |
| **SC29-1_b** | 7 | 4 | 3 | 18 | 17 | 0 | 22 | 25 | 0 | 51 | 41 | 28 | 0 | 7 | 31 |
| **SC29-8_b** | 7 | 4 | 3 | 18 | 15 | 0 | 22 | 25 | 0 | 53 | 44 | 27 | 0 | 7 | 31 |
| **SW21-5_b** | 8 | 2 | 1 | 13 | 13 | 0 | 21 | 25 | 0 | 57 | 53 | 30 | 0 | 4 | 40 |
| **T103-1_B** | 11 | 2 | 4 | 16 | 16 | 0 | 20 | 28 | 0 | 34 | 59 | 23 | 0 | 7 | 37 |
| **T126-1_N** | 11 | 2 | 6 | 15 | 13 | 0 | 17 | 24 | 0 | 42 | 57 | 25 | 0 | 8 | 30 |
| **T128-1_B** | 4 | 1 | 6 | 13 | 15 | 0 | 20 | 31 | 0 | 44 | 53 | 22 | 0 | 3 | 45 |
| **Tptr3-1_B** | 11 | 1 | 5 | 13 | 13 | 0 | 20 | 27 | 0 | 44 | 54 | 24 | 0 | 8 | 36 |
