## Supplemental Material for "The virulence gene *ToxB* is both amplified and disrupted by transposons in the wheat pathogen *Pyrenophora tritici-repentis*"

Article acceptance date: Click here to enter a date.

The following Supporting Information is available for this article:

**Fig. S1** Zoomed in LASTZ alignment of select isolates T126-1 (track 1; tig206), K11 (track 2; tig47), G9-7 (track 3; tig154), and Den17 (track 4; tig96) which lack *ToxB/toxb* with ToxB containing (green triangels) isolate I-73-1 *ToxB* multi-copy contig12 (black bar). Variable gap sizes can be seen in the *ToxB* negative isolates, ranging from ~50 Kb to ~375 Kb. Gap sizes alter slightly when compared the to single copy carrier Tptr3-1 or *toxb* carrier D308 (not shown).

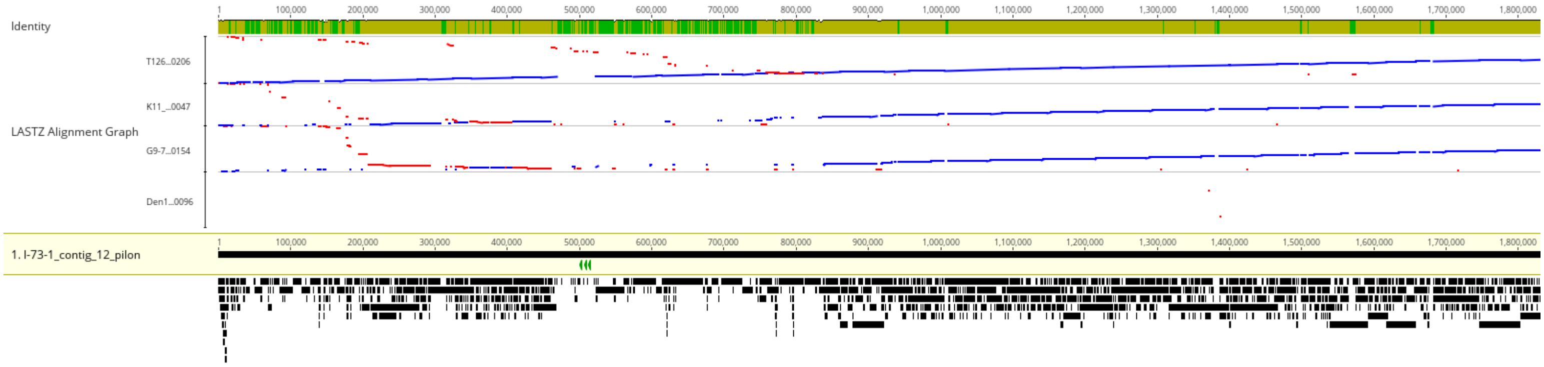


**Fig. S2** Dotplots showing single reads (y-axes) covering all copies of *ToxB* in the assemblies of four multicopy isolates (x-axes).

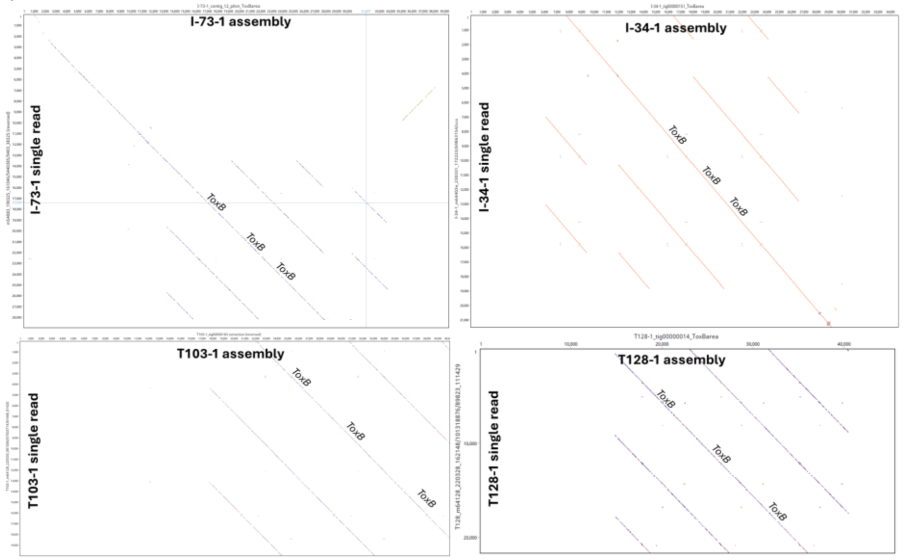

**Fig. S3** Read coverage around *ToxB* in isolate Alg3-24 **a** Read coverage of *ToxB* region in tig35 (Chr05) in isolate Alg3-24; **b** Read coverage of multi *ToxB* copy tig112 (Chr10) in isolate Alg3-24. A drop in coverage is visible for the second copy.

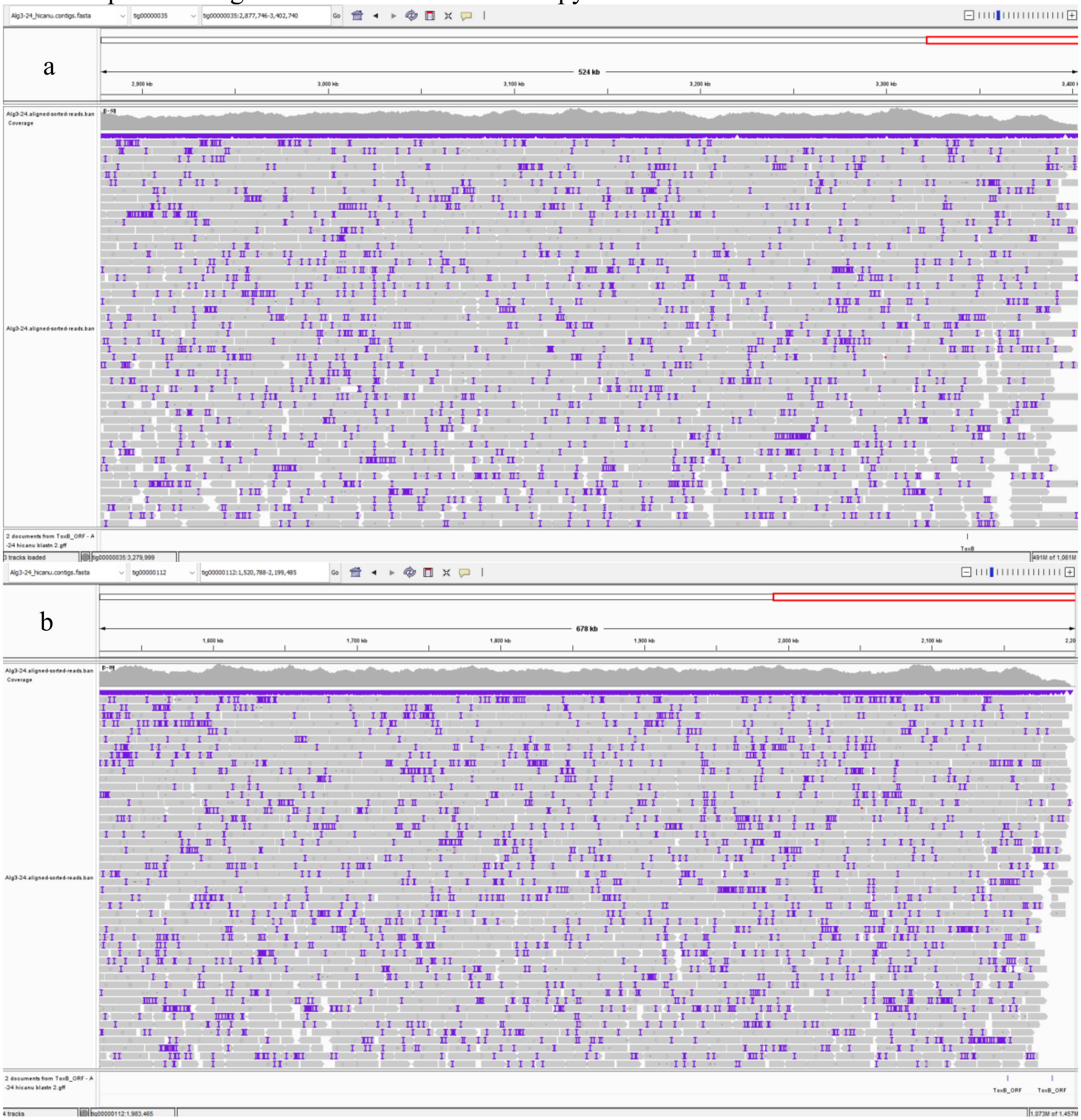

**Fig. S4** Example of self-alignments done with isolate T128-1 which were used to locate edges of the *ToxB* replicative unit. **a** linear dotplot showing the unidirectional nature of the duplications and helps locate the specific start and end locations; **b** circular alignment of the same 45 kbp shows the replication of the *ToxB* region more clearly, with duplications separated by a sequence junction. The alignments also revealed a pocket of nearby putative transposons and that the entire region is flanked by a hAT-like transposon.

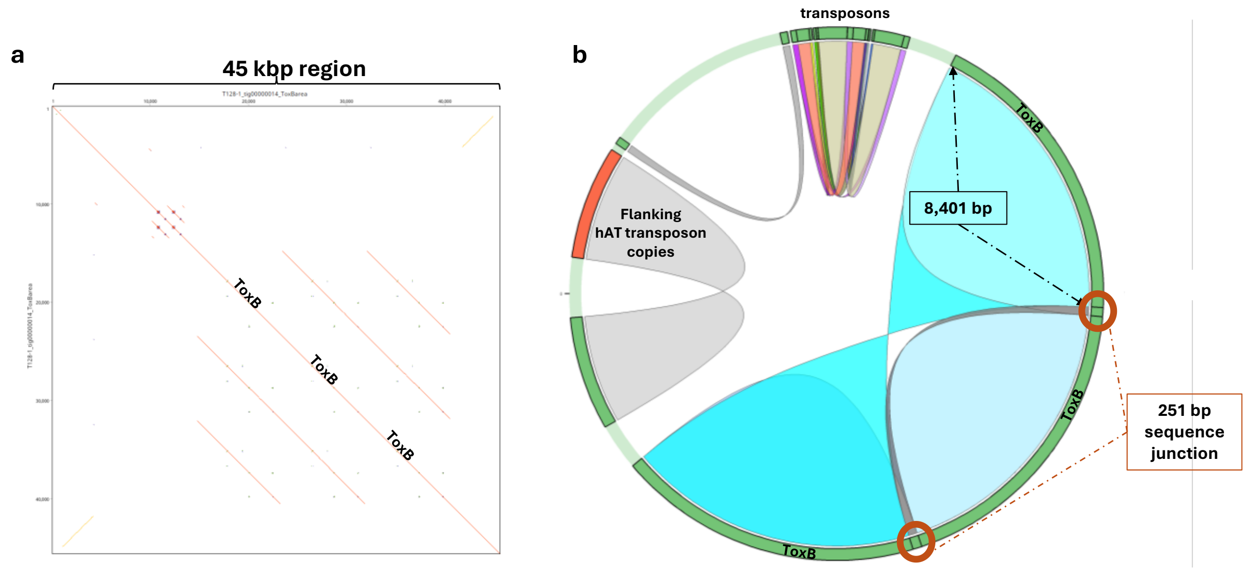

**Fig. S5** Linear alignments of the *ToxB/toxb* regions of Chr04 in *Pyrenophora tritici-repentis*. Colored arrows along the chromosome (black line) show *ToxB* (yellow), *toxb* (white), ORFs (green), transposons (reds and oranges), and the sequence junction between duplicated regions (black). Due to the alignment scale, vertical yellow and black arrows have been added to aid in visualizing the positions of *ToxB* and the sequence junction respectively. The linking blocks drawn between isolates show aligned DNA identity. Alignments less than 500 bp were omitted for clarity, as were most alignments between copies of the *hAT-1_Ptr* transposon. Region duplications in isolates I-73-1 and AlgH1 are apparent when compared to the single *ToxB* containing isolate Tptr3-1. The position and number of *hAT-1_Ptr* transposons in the multi-copy isolates T128-1, I-73-1, and AlgH1 combined with the conserved edges suggest independent duplication events via the same mechanism. *ToxB* disruption via Copia retroelements is also visible in isolates D308 (*Copia-1_Ptr*) and SC29-8 (*Copia-2_Ptr*). Unedited version.

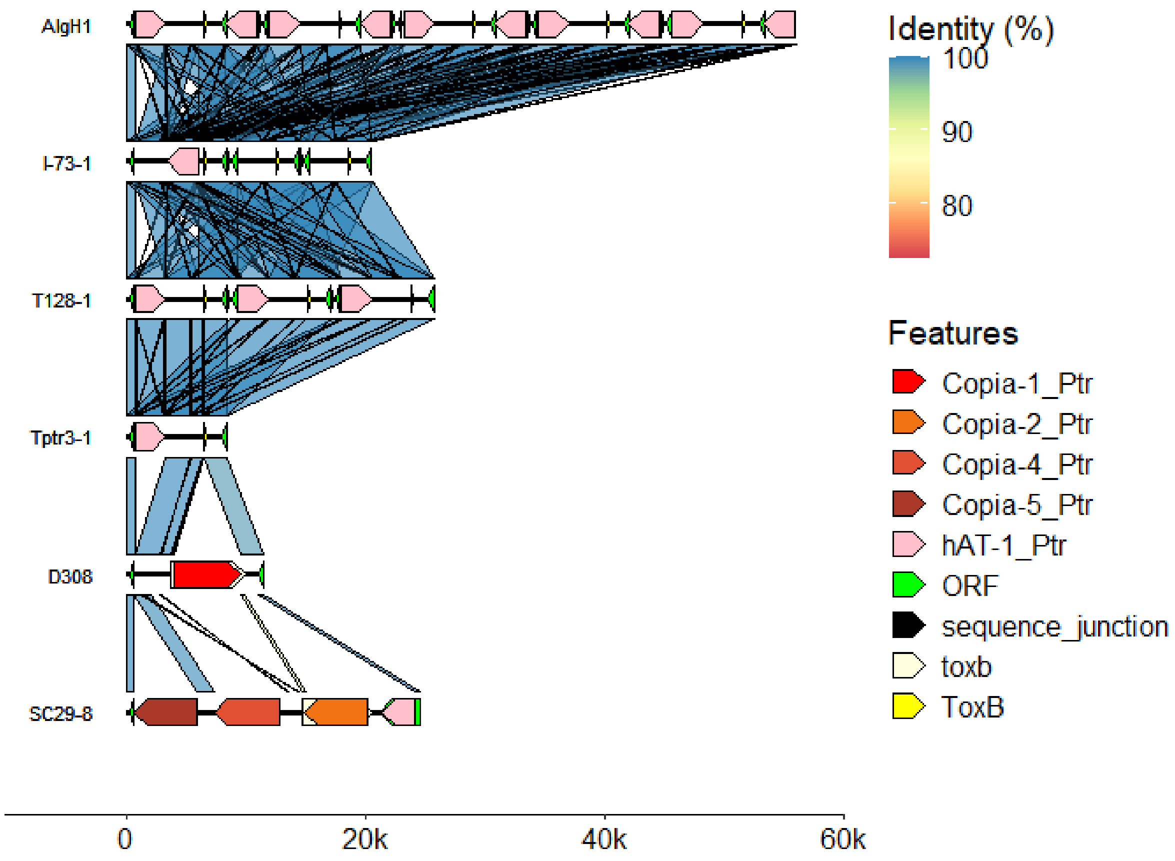

**Fig. S6** Density plots showing the proportion of 50 kbp genome segments with the number of TE annotations shown on the x-axis. The density plots is shown for isolate I-73-1 using two different TE detection methods, EDTA (top graph) and EarlGrey (bottom graph). The larger blue plot at the top shows the genome wide TE density, while the smaller graphs below show the TE density for each chromosome individually. Bi-modal peaks show chromosomes that have regions with a higher TE density.

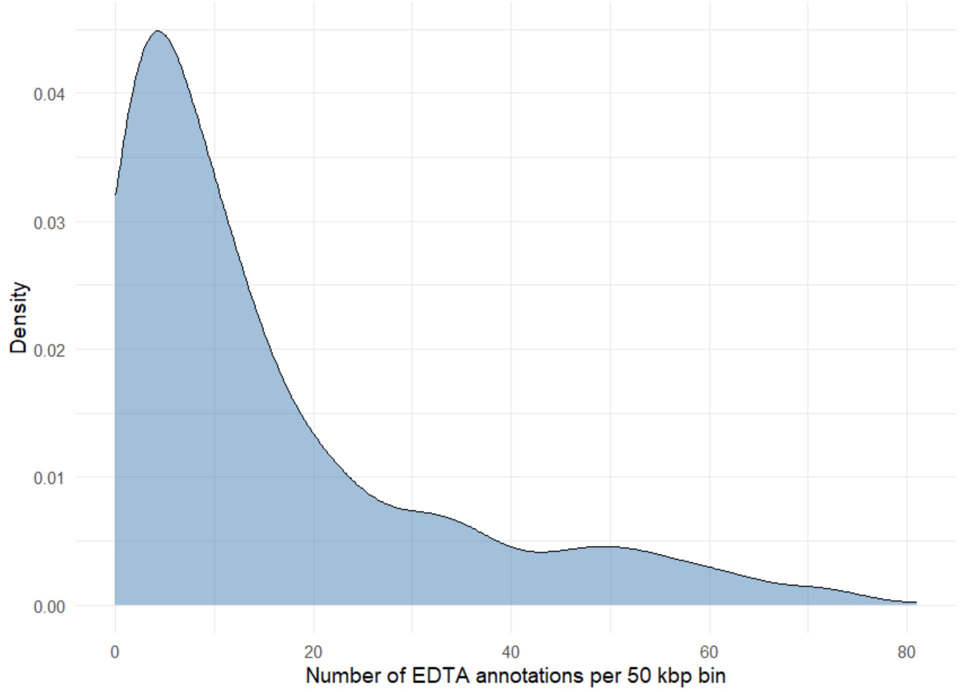


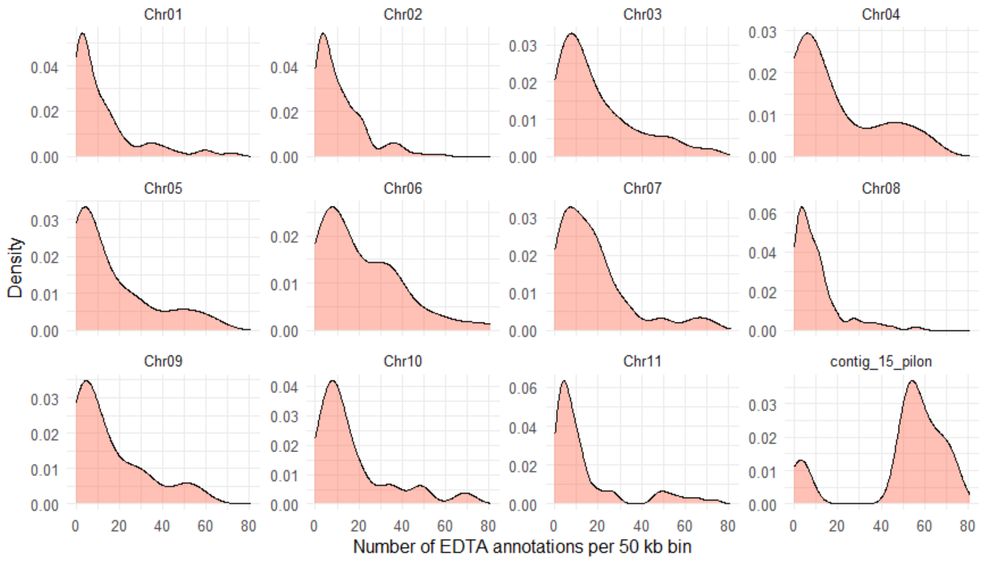

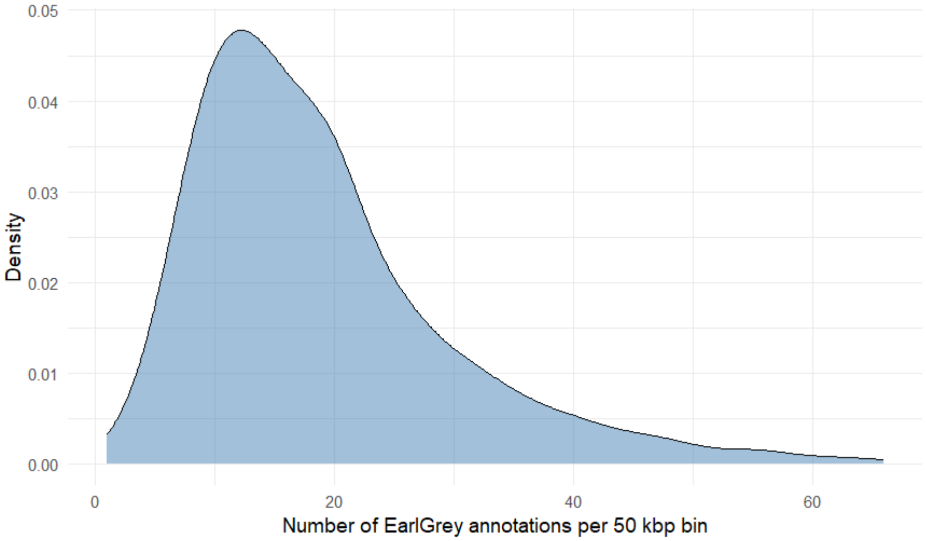
Density
Number of EarlGrey annotations per 50 kbp bin

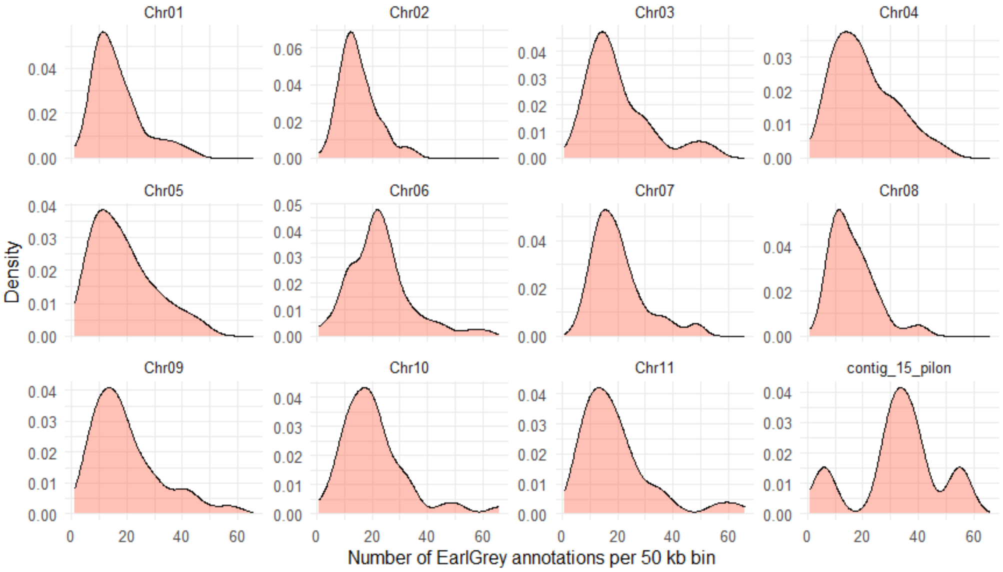
Chr01
Chr02
Chr03
Chr04
Chr05
Chr06
Chr07
Chr08
Chr09
Chr10
Chr11
contig_15_pilon
Density
Number of EarlGrey annotations per 50 kb bin

**Fig. S7** Dotplot alignment (30 bp sliding window) of elements Copia-1_Ptr from D308 (x-axis) and *Copia-2_Ptr* from SC29-1 (y-axis) which were inserted in *ToxB* ORF of the respective isolates. Pairwise alignment with MUSCLE showed an identity of ~51%.

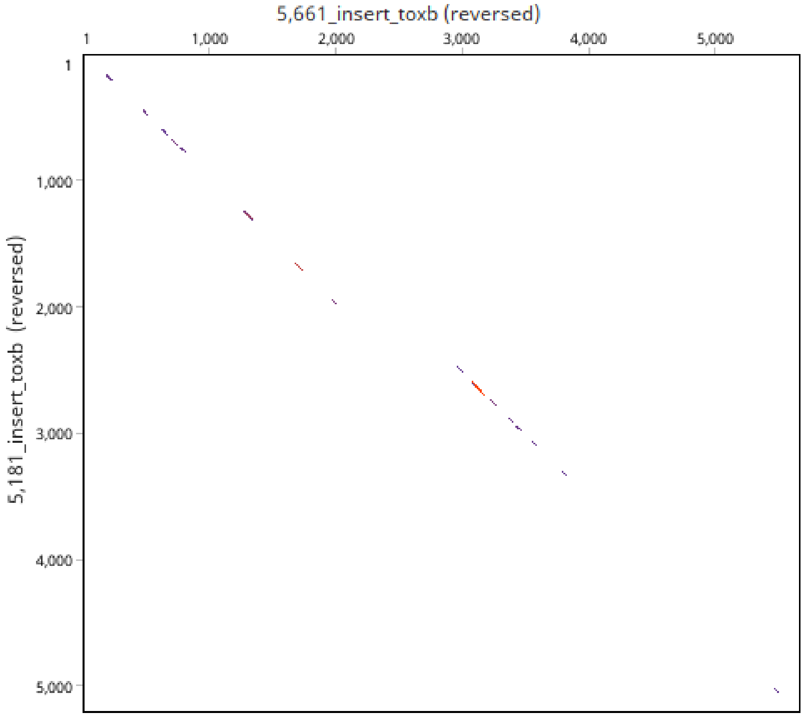


**Fig. S8** Schematic of a histone replicating hAT transposon present near *ToxB* in *Pyrenophora tritici-repentis* isolate T128-1. Target site duplications (green) flanking terminal inverted repeats (white) mark the transposon edges when compared with the non-transposon containing intergenic sequence (grey) of isolate I-73-1. Open-reading frames within the transposon are marked by yellow arrows which show orientation. Figure is not to scale.

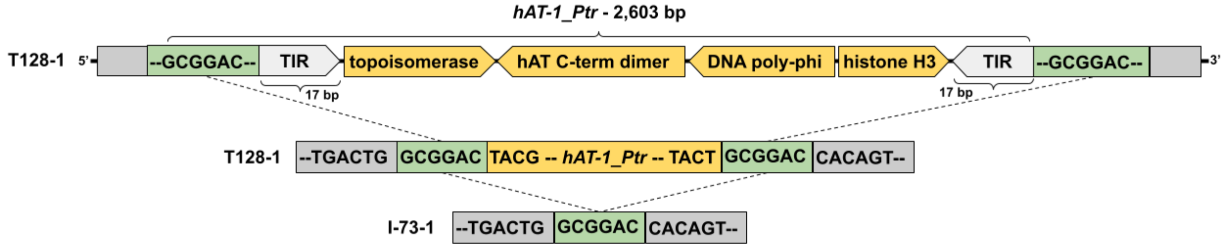

**Fig. S9** Maximum likelihood tree (50 bootstraps) of transposable elements annotated with EDTA and/or EarlyGrey which overlapped with BLASTN searches of the 251 bp sequence-junction. Red highlights are those residing within the duplication region. A limited representative sampling of isolates were used: I-73-1, T128-1, Tptr3-1, SC29-8, and D308.

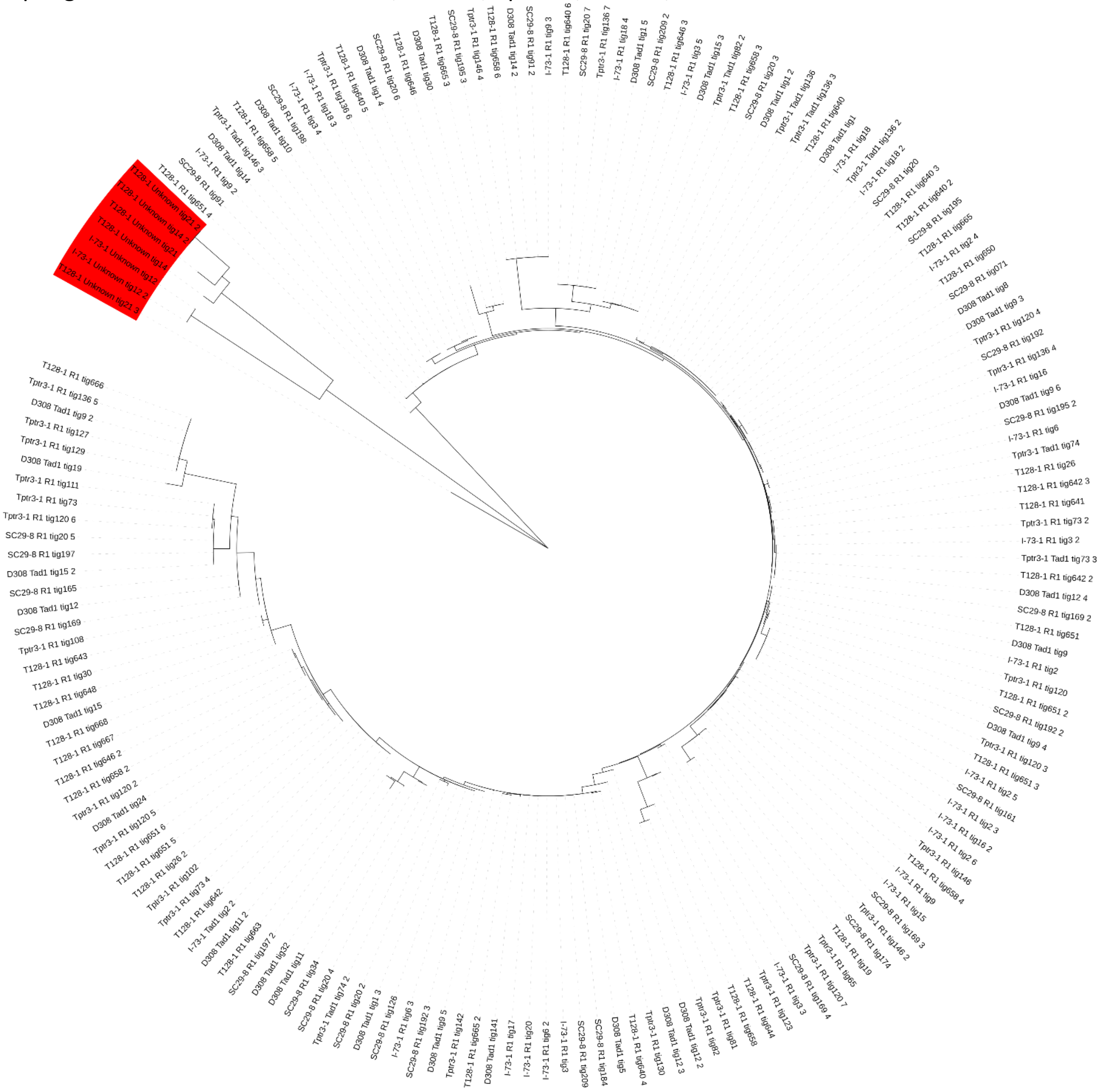

**Fig. S10** Generalized schematics of known Helitrons elements. Canonical Helitron1 adapted from Kaptinov and Jurka (2007); Helitron2 adapted from Boa and Jurka (2013); Helentron adapted from Thomas and Pritham (2015); and *Fusarium* Helitrons adapted from Chellapan et al. (2016). RepHel element may be present in all Helitrons however the majority of documented Helitrons lack these domains so are omitted from all but the canonical Helitron1. Loops and asymmetrical terminal inverted repeats (arrows) are shown along with the left-terminal sequences and right-terminal sequences where R = A or G.

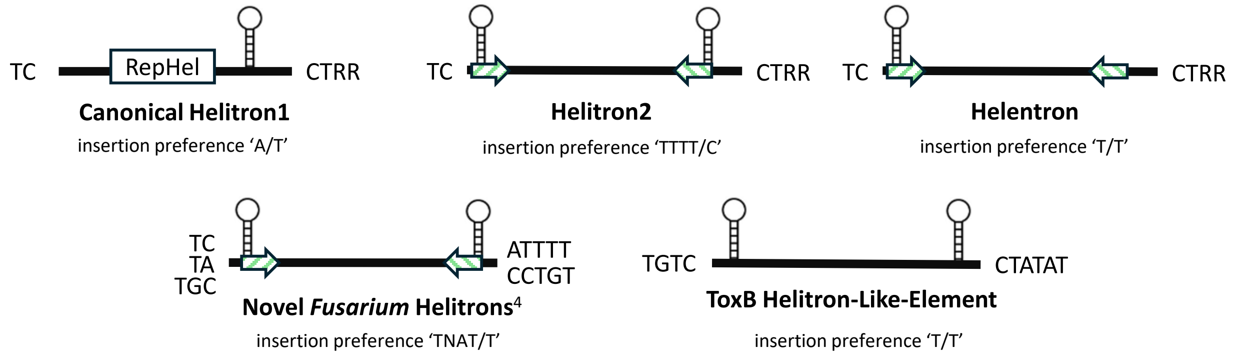


**Fig. S11** Loop structures found near the terminal edges of *FoHeli1* in *Fusarium oxysporum* and *ToxB-HLE* in *Pyrenophora tritici-repentis*; **a** 5' LTS loop of FoHeli1; **b** 3' loop of *FoHeli1* beginning 51 bp from the RTS; **c** 5' LTS loop of ToxB-HLE (red text = 'TGTC' start sequence); **d** 3' loop of *ToxB-HLE* beginning 72 bp from RTS.

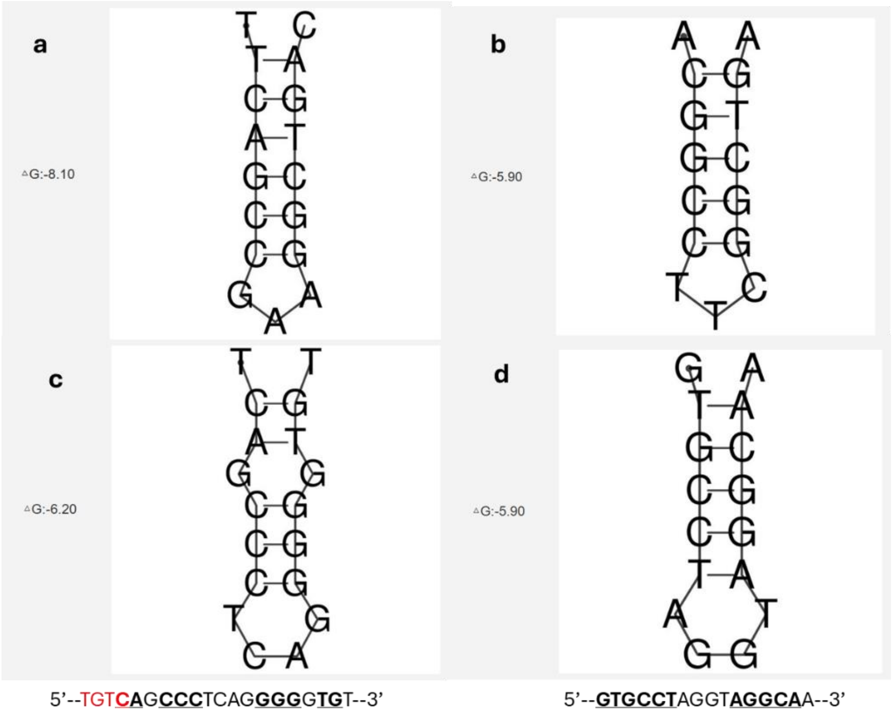

**Fig. S12** Geneious annotations surrounding copies of *ToxB* (green arrows) in isolate 92-171R5. The duplication of *ToxB* and surrounding sequences are marked in light blue. All other annotations are transposons, or transposon remnants annotated by EDTA. The duplicated *ToxB* is flanked on either side by undefined retrotransposons.

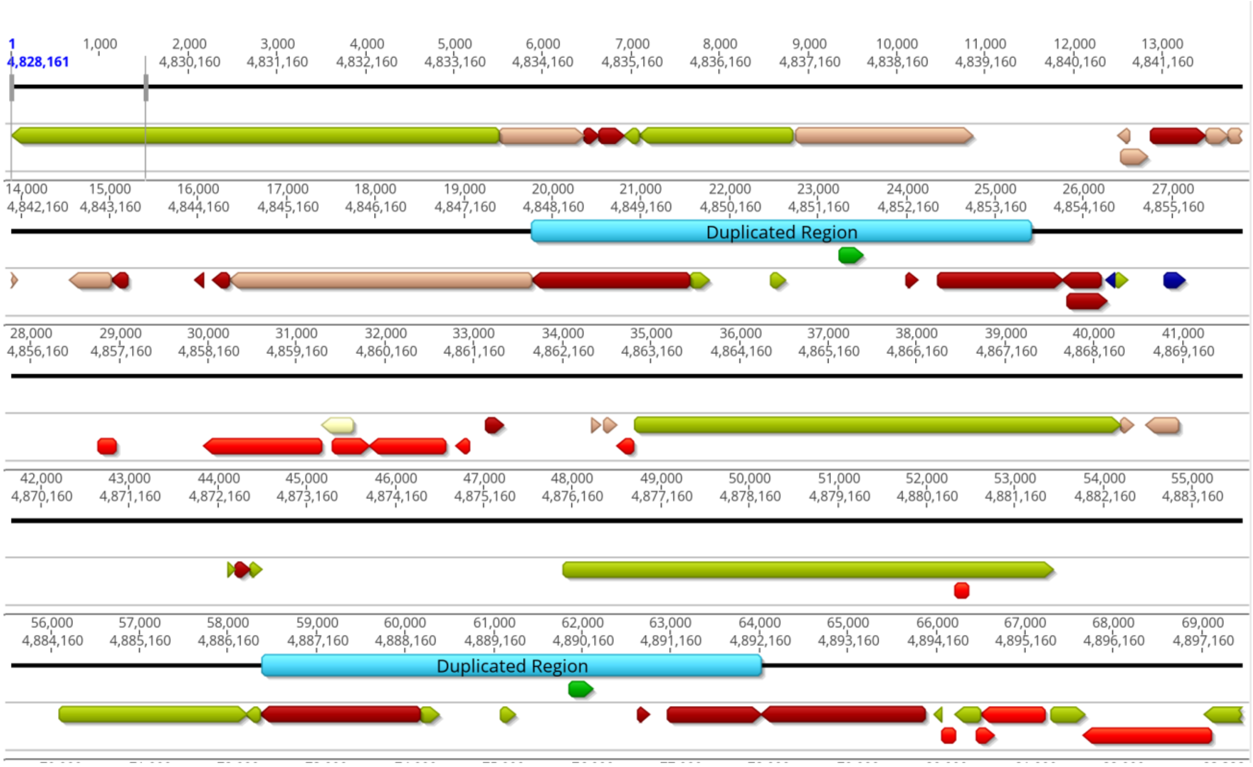

**Table S1** Details of transposons and insertions near or within the *ToxB* ORF in *Pyrenophora tritici-repentis*.

| BankIt | Accession | Length | Name | Category | LTR/TIR length (bp) |
|---|---|---|---|---|---|
| 2984043 | PV975795 | 5,661 | *Copia-1_Ptr* | LTR retrotransposon | 192 LTR |
| 2984043 | PV975796 | 5,181 | *Copia-2_Ptr* | LTR retrotransposon | 165 LTR |
| 2984043 | PV975797 | 5,549 | *Copia-3_Ptr* | LTR retrotransposon | 195 LTR |
| 2984043 | PV975798 | 5,396 | *Copia-4_Ptr* | LTR retrotransposon | 240 LTR |
| 2984043 | PV975799 | 5,251 | *Copia-5_Ptr* | LTR retrotransposon | 172 LTR |
| 2984043 | PV975800 | 5,345 | *Copia-6_Ptr* | LTR retrotransposon | 102 LTR |
| 2984043 | PV975801 | 6,070 | *Copia-7_Ptr* | LTR retrotransposon | 202 LTR |
| 2984043 | PV975802 | 2,603 | *hAT-1_Ptr** | DNA transposon | 17 TIR |
| | - | 2,805 | *hAT-1.2_Ptr*** | DNA transposon | 17 TIR |
| 2984043 | PV975803 | 4,036 | *hAT-2_Ptr* | hAT DNA transposon | 257 LTR |
| 2984043 | PV975804 | 1,060 | *hAT-3_Ptr* | DNA transposon | - |
| 2984043 | PV975805 | 3,005 | *hAT-4_Ptr* | DNA transposon | 20 TIR |
| 2984043 | PV975806 | 2,110 | *Pogo-1_Ptr* | DNA transposon | 25 TIR |
| 2984043 | PV975807 | 5,620 | *Ty3-1_Ptr* | LTR retrotransposon | 256 LTR |
| 2984043 | PV975808 | 6,770 | *Ty3-2_Ptr* | LTR retrotransposon | 454 LTR |
| | - | 5,601 | - | Possible non-LTR retrotransposon | - |
| | - | 1,677 | - | LTR transposon | 147 LTR |
| | - | 240 | - | Abandoned LTR from Copia-4 | - |
| | - | 202 | - | Abandoned LTR from Copia-7 | - |
| | - | 170 | - | Abandoned LTR from unknown | - |

*Defined in Hafez et al., 2023

***hAT-1_Ptr* with the 202 abandoned LTR from *Copia-7_Ptr* inserted within

**Table S2** Counts of transposons and the sequence junction throughout the genomes of P.tritici-repentis. Counts based on BLASTN 90% identity over 90% of the sequence. [SEE SEPARATE FILE]

**Methods S1** Sequencing details of genomic DNA from Pyrenophora tritici-repentis.

The gDNA (1.7-4.8µg) was sheared using a MegaRuptor 3 system (Diagenode) to a target size of 12-16kb prior to library preparation. Sheared DNA was prepared for sequencing following the PacBio recommended procedure (PN 102-166-600 APR2022). Briefly, the DNA is treated for removal of single-stranded overhangs, damage repaired, end prepared, and ligated to a SMRTbell adapter with the SMRTbell Prep kit 3.0 (REF: 102-141-700). To target recovery of molecules ≥10 kb, SMRTbell libraries were size selected on the BluePippin system (Sage Science) using the 0.75% Agarose Dye-Free Gel Cassette (BLF7510) with the S1 Marker. Further libraries were quantified with Qubit (Thermo Fisher Scientific) and size distribution was estimated with FEMTO Pulse (Agilent Technologies) instruments. Finally, multiplexed libraries were bound to the sequencing polymerase enzyme using the Sequel II Binding Kit (REF:102-194-100) before loading at an on-plate loading concentration (OPLC) of 90pM on the PacBio Sequel IIe System (Pacific Biosciences) using the Sequel II Sequencing Kit (REF: 101-820-200) with a SMRT Cell 8M Tray (REF: 101-389-001). Sequence data was collected for a 30h movie acquisition times preceded with a 2h pre-extension time and 2h of adaptive loading. Subreads were generated using the SMRT Link v.11.1.0.166339; Chemistry Bundle: 11.1.0.154383; Params: 11.1.0; and SMRT Link v.12.0.0.177059; Chemistry Bundle: 12.0.0.172289; Params: 12.0.0.
